# Population structure of Apodemus flavicollis and comparison to Apodemus sylvaticus in northern Poland based on RAD-seq

**DOI:** 10.1101/625848

**Authors:** Maria Luisa Martin Cerezo, Marek Kucka, Karol Zub, Yingguang Frank Chan, Jarosław Bryk

## Abstract

**Background:** Mice of the genus *Apodemus* are one the most common mammals in the Palaearctic region. Despite their broad range and long history of ecological observations, there are no whole-genome data available for *Apodemus*, hindering our ability to further exploit the genus in evolutionary and ecological genomics context.

**Results:** Here we present results from the double-digest restriction site-associated DNA sequencing (ddRAD-seq) on 72 individuals of *A. flavicollis* and 10 *A. sylvaticus* from four populations, sampled across 500 km distance in northern Poland. Our data present clear genetic divergence of the two species, with average p-distance, based on 21377 common loci, of 1.51% and a mutation rate of 0.0011 - 0.0019 substitutions per site per million years. We provide a catalogue of 117 highly divergent loci that enable genetic differentiation of the two species in Poland and to a large degree of 20 unrelated samples from several European countries and Tunisia. We also show evidence of admixture between the three *A. flavicollis* populations but demonstrate that they have negligible average population structure, with largest pairwise F_ST_ < 0.086.

**Conclusion:** Our study demonstrates the feasibility of ddRAD-seq in *Apodemus* and provides the first insights into the population genomics of the species.

## Background

Mice of the genus *Apodemus* (Kaup, 1829) (Rodentia: Muridae) are one the most common mammals in the Palaearctic region [38]. The genus comprises of three sub-genera (*Sylvaemus, Apodemus* and *Karstomys*) [38], however the systematic classification of the 20 species belonging to the genus [17] is not fully settled [32]. In the Western Palearctic, the yellow-necked mice *A. flavicollis* (Melchior, 1934) and the woodmice *A. sylvaticus* (Linnaeus, 1758) are widespread, sympatric and occasionally syntopic species. They are often difficult to distinguish morphologically in their southern range [27], but in the Central and Northern Europe both are easily recognisable by the full yellow collar around the neck of *A. flavicollis*, which only forms a narrow elongated spot on the breast in *A. sylvaticus* [51].

Their prevalence in Western Palearctic and common status in Western and Central Europe made them one of the model organisms to study post-glacial movement of mammals [21, 40]. Both species have traditionally been studied in a parasitological context, as one of the vectors of *Borellia*-carrying ticks *Ixodes ricinus*, who often feed on *Apodemus* [42, 57], tick-borne encephalitis virus [14] and hantaviruses [30, 45] and have been used as markers for environmental quality [35, 62]. Lastly, they have extra-autosomal chromosomes, called B chromosomes, with varied distribution among the populations [55] and suggested involvement in a variety of physiological phenomena, from cell division and development to immune response [63].

Previous studies on *Apodemus* typically employed a small number of microsatellite [58] and mtDNA markers [21, 37, 39, 40], which are insufficient to learn about the species’ population structure and admixture patterns in detail, or to identify loci under selection. In the absence of high-quality reference genome, which remains cost-prohibitive for complex genomes, whole-genome marker discovery enabled by restriction site-associated DNA sequencing presents a cost-effective method to study species on a population scale even with no previous genetic and genomic resources available [5].

Here we employ the double-digest restriction site-associated DNA sequencing (ddRAD-seq) to elucidate the genetic structure and connectivity of three populations of *A. flavicollis* and compare it to a population of *A. sylvaticus* in Poland. We demonstrate clear divergence between the two species and very low differentiation between populations of *A. flavicollis*. Our results provide the first estimates of population parameters in *A. flavicollis* based on thousands of loci, calculation of p-distance between the two *Apodemus* species, as well as a selection of loci enabling their accurate identification.

## Results

### Sequencing and variant calling

The sequencing produced a total of 92741120 reads. The number of reads per individual varied from 346810 to 4157586, with an average of 1078385 reads per individual and median of 905786,5 (Supplementary Table S2). The best parameters for calling the stacks and variants for the entire dataset were: minimum number of identical, raw reads required to create a stack m = 2, number of mismatches allowed between loci for each individual M = 4 and number of mismatches allowed between loci when building the catalogue n = 5 (Supplementary Figure S1). The best parameters calculated for *A. flavicollis* samples only were: m = 2, M = 4 and n = 3 (Supplementary Figure S3). The coverage per sample ranged from 4.95x to 26.20x with an average of 10.13x and median of 9.32x for the entire dataset (Supplementary Figures S2 and S4).

### SNPs and loci co-identification rates

Analysis of the duplicated samples showed that loci and allele misassignment rates were of similar magnitude, on average, between all pairs of duplicates. The duplicate pair F06-B02 showed the highest discrepancy between loci, of 10 %, and also between alleles, of 8 %. When only shared loci were included in the comparisons, all four sets of duplicates showed on average 0.5% ± 0.2% SNPs called differently (Table 1).

**Table 1:**
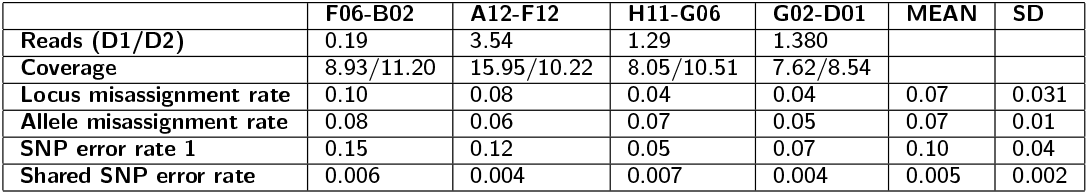
Error rates calculated by comparing four sets of duplicated samples. D1/D2: ratio of reads from Duplicate 1 to Duplicate 2. Locus misassignment rate: the percentage of unidentified loci, calculated by dividing the number of loci found only in one of the duplicates by the total number of loci in each sample. Allele misassignment rate: the percentage of mismmatches between the IUPAC consensus sequences between homologous loci from each pair of duplicates. SNP error rate 1: the percentage of different SNPs called in each of the duplicated samples using either 10178 SNPs. Shared SNP error rate: the percentage of different SNPs called in each of the duplicated samples after excluding missing data between duplicate samples

### Comparison of *A. flavicollis* and *A. sylvaticus*

The number of assembled loci per individual ranged from 46286 to 117366 (mean: 73711, median: 71395, standard deviation: 29917). 52494 loci passed the population filters established for species differentiation (see Methods, section “Variant calling and filtering”), representing 8,3% of the total 632063 loci included in the catalogue. Out of 158144 SNPs called, 60366 (38.1%) were removed after filtering for minor allele frequency (MAF) and 52298 (33%) were removed after failing the HWE test at p<0.05; further 35302 (22.3%) were removed due to a minimum mean depth lower than 20, leaving 10178 SNPs (6.6%) to be used in the downstream analyses (Figure 1). PCA plot of the first two components (Figure 2), accounting for 13.13% of the total variance, shows differentiation of the two species but also distinguish different populations of *A. flavicollis*.

**Figure 1:**
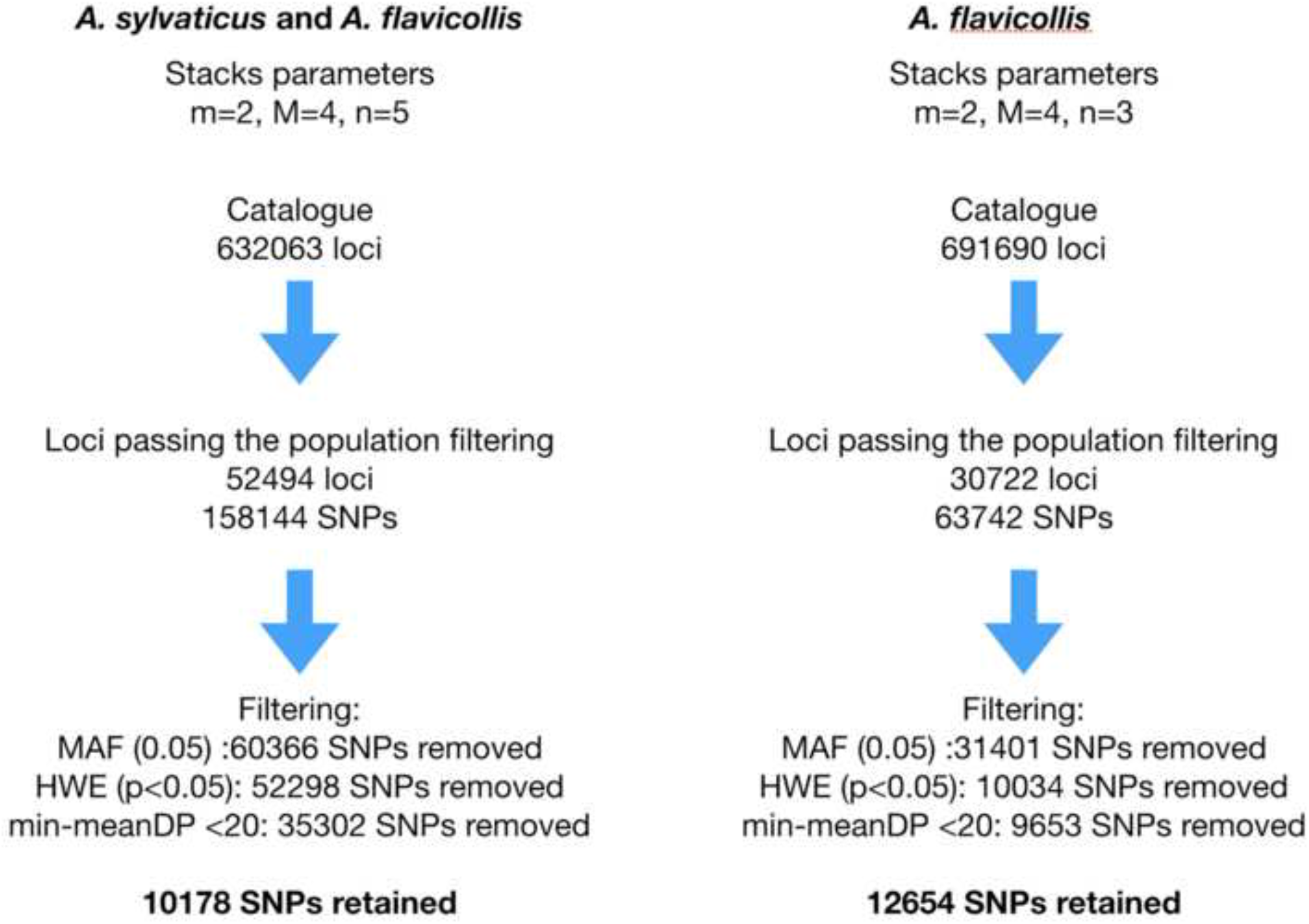
Summary of cataloque construction and SNP filtering steps for the complete dataset (left) and *Apodemus flavicollis* dataset. The graphic includes: Stacks parameters values (m, M, n), number of loci in the catalogue, number of SNPs filtered by minor allele frequency (MAF), which failed the Hardy-Weinberg equilibrium test at p<0.05 (HWE), SNPs removed due to an average depth, across individuals, lower than 20 (min-meanDP) and the total number of SNPs retained for further analysis

**Figure 2:**
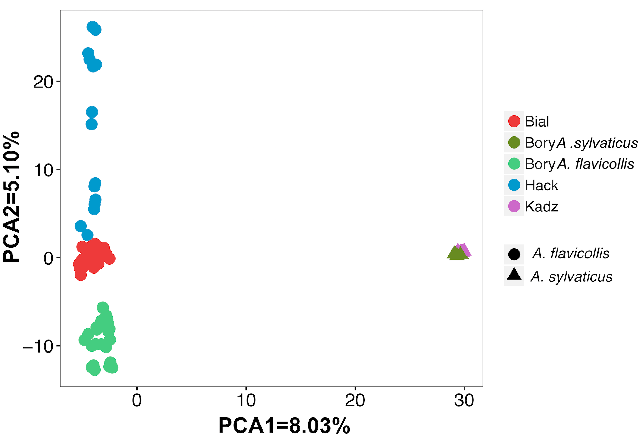
Principal Component Analysis of all samples analysed in the study. Each point represents one sample; the shape of the point represents the species (circles: *Apodemus flavicollis* (n = 72), triangles: *Apodemus sylvaticus* (n = 10), whereas the colour represents the location where the samples were collected: Bial - Biaowiea, Kadz - Kadzido, Hack - Haki, Bory - Bory Tucholskie.

Similarly, the phylogenetic tree shows *A. sylvaticus* as a separate clade to the three populations of *A. flavicollis*, with *A. flavicollis* from geographically closer regions (Biaowiea and Haki, 50 km) grouped closer than a population from Bory Tucholskie, 450 km away from Biaowiea (Figure 3). The *A. sylvaticus* and *A. flavicollis* clusters have high bootstrap value support (100% and 99% respectively).

**Figure 3:**
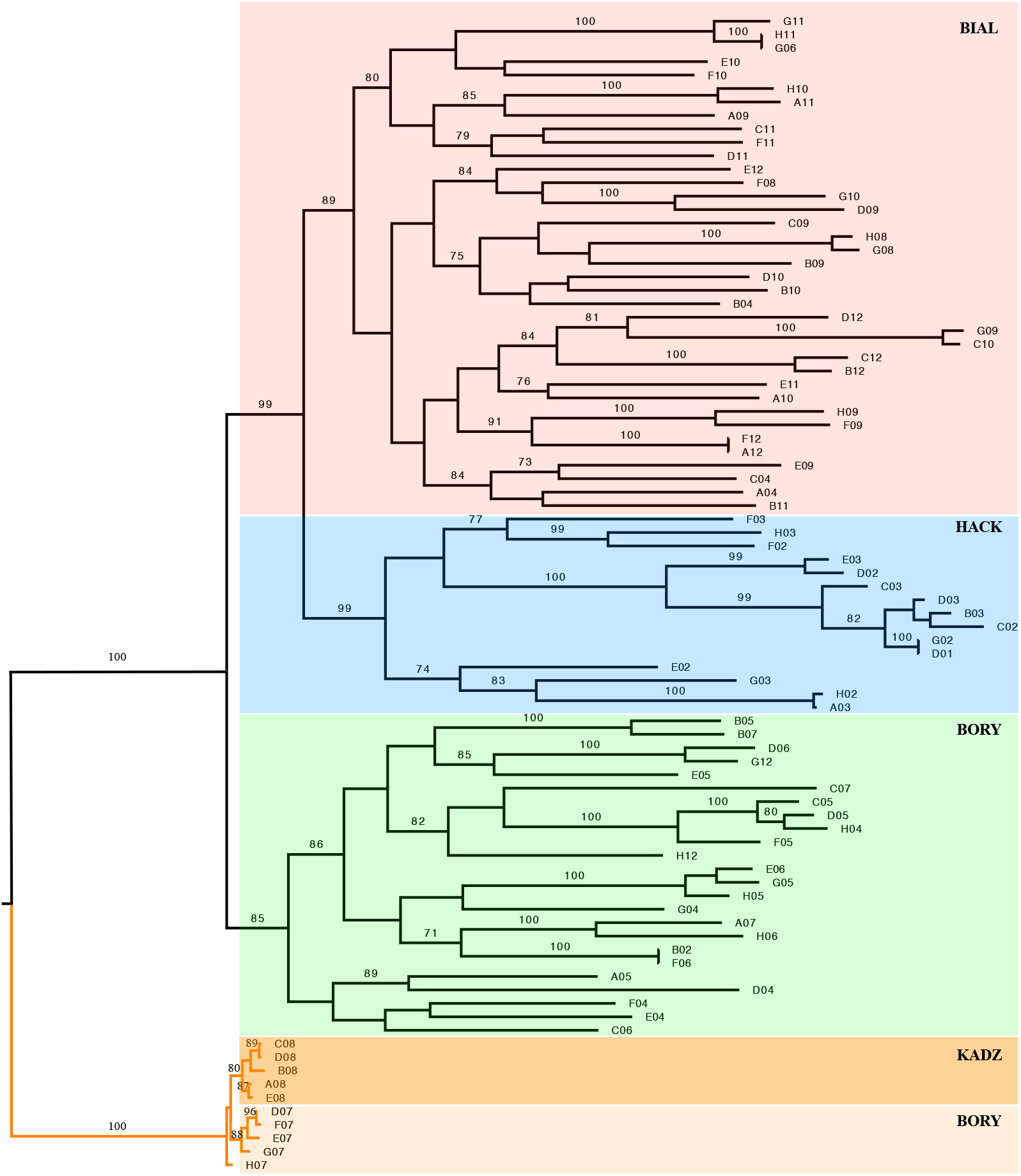
Maximum likelihood phylogenetic tree of all the samples analysed in the study. Colour represents the species: *A. sylvaticus* (n=10) in orange and *A. flavicollis* (n=72) in black. Duplicates samples are included: F06-B02 from Bory Tucholskie, F12-A12 and H11-G06 from Bialowieza and G02-D01 from Hacki. Bootstrap support values from 100 replicates are indicated at the nodes of the tree. Bial - Biaowiea, Kadz - Kadzido, Hack - Haki, Bory - Bory Tucholskie.

We then investigated the suitability of the loci we identified on Polish populations to distinguish *A. sylvaticus* and *A. flavicollis* from other European populations. The genotyping of the extra 10 samples from each species (see Methods) produced 179763 SNPs. 62158 (34.58%) were removed after filtering for MAF and 69125 (38.45%) were removed after failing the HWE test at p<0.05; further 42054 (23.39%) were removed due to a minimum mean depth lower than 20 and 5203 (2.89%) were removed due to more than 5% missing data, leaving 1223 SNPs (0.68%) to be used in the downstream analyses.

The first axis of the PCA plot (Figure 4) constructed from this data accounts for the 65.73 % of the total variance and shows clear differentiation between the two species. All the *A. flavicollis* samples cluster with the Polish *A. flavicollis* samples, while all but Tunisian samples of *A. sylvaticus* cluster with the Polish samples of the same species. Tunisian *A. sylvaticus* appear as a separate cluster but still closer to the *A. sylvaticus* group. The catalogue of loci used for species identification is included in the Supplementary Materials, Section 6.

**Figure 4:**
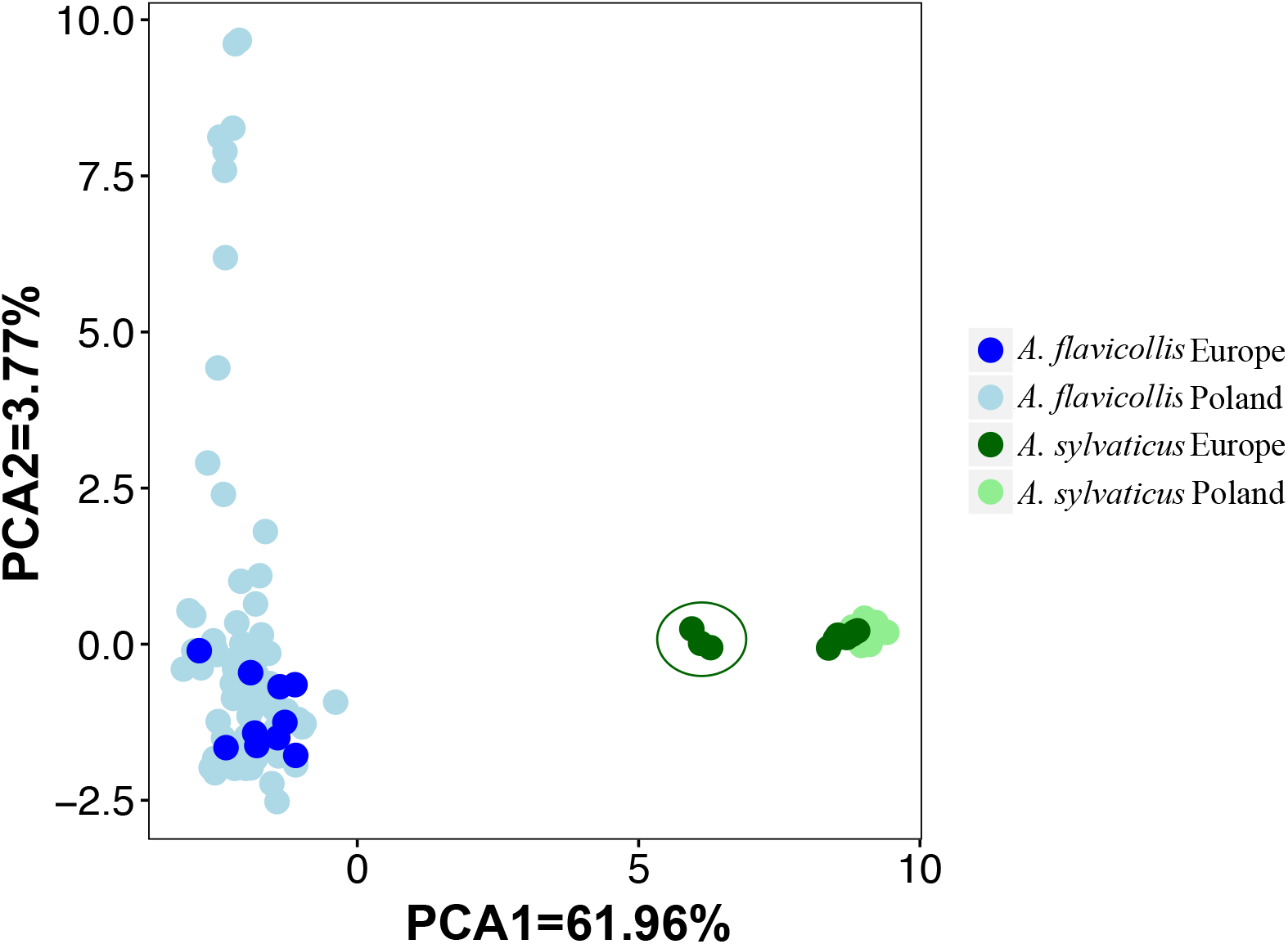
Species identification through Principal Component Analysis using a catalogue of 632060 loci and 1223 final SNPs. Light colours represent samples from Poland while dark colours represent samples from other European regions and Tunisia (collectively named “Europe”; Tunisian samples are marked with a circle). Green: *A. sylvaticus*, blue: *A. flavicollis*.

### Genetic diversity and population structure of *A. flavicollis*

The number of assembled loci per individual in the Polish populations ranged from 46286 to 117366 (mean: 72738, median: 70592, stdev: 12575). 30722 loci passed the population filters established for population differentiation, representing and 4,43% of the total 691960 loci included in the catalog. Out of 63742 SNPs called, 31401 (49.26%) were removed after filtering for MAF and 10034 (15.74%) were removed after failing the HWE test at p<0.05. Further 9653 (15.14%) were removed due to a minimum mean depth lower than 20, leaving 12654 (19.85%) SNPs to be used in the downstream analyses (Figure 1).

PCA plot (Figure 5) shows differentiation between the three Polish *A. flavicollis* populations, with PC1 and PC2 cumulatively explaining 10.47% of the total variance. Haki population shows larger diversity than the other populations, with some Haki individuals closer to Biaowiea individuals than to others from this location. Phylogenetic tree (Figure 6) supports this pattern of differentiation. Bory Tucholskie and Haki populations each form a cluster with a 100% of bootstrap support value, whereas Biaowiea forms a third cluster with an 95% of bootstrap support. Biaowiea and Bory Tucholskie population together form a large cluster with a 100% bootstrap support.

**Figure 5:**
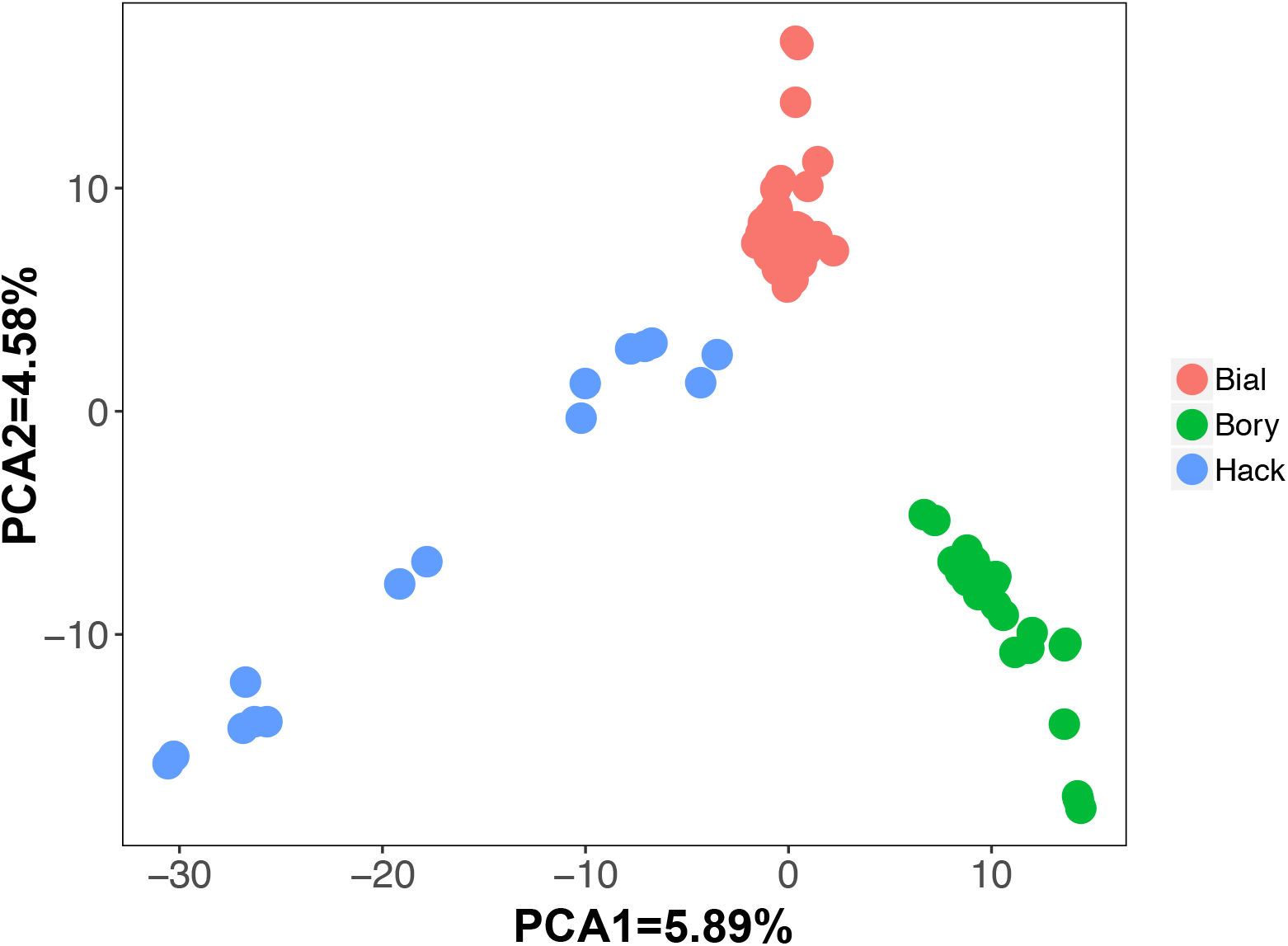
PCA plot showing Polish samples of *A. flavicollis* from Biaowiea (red) (n=35), Haki (blue) (n=14) and Bory Tucholskie (green) (n=23). Bial - Biaowiea, Kadz - Kadzido, Hack - Haki.

**Figure 6:**
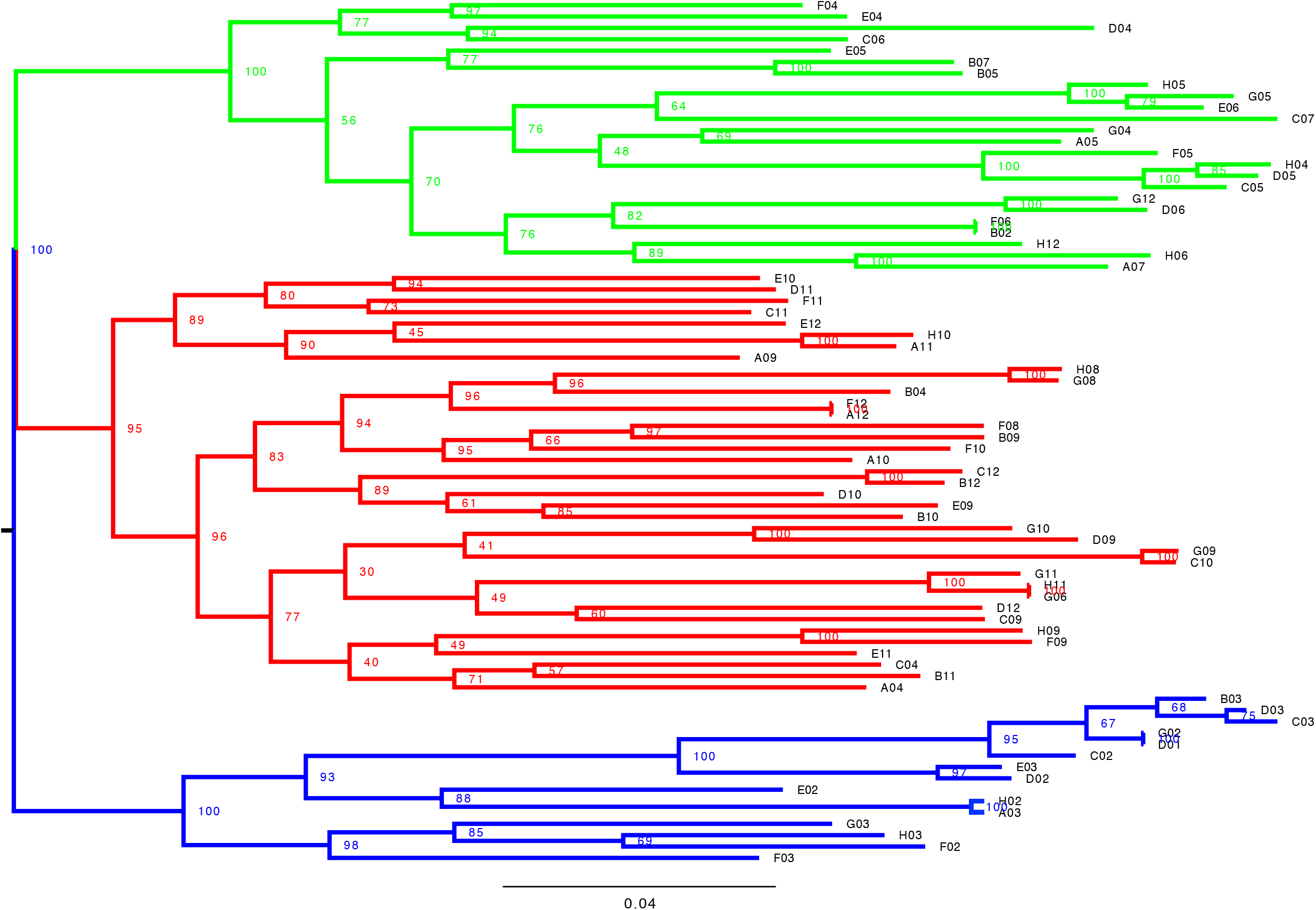
Maximum ilkelihood phylogenetic tree of n = 72 *A. flavicollis* samples from Biaowiea (red, n = 35), Haki (blue, n = 14) and Bory Tucholskie (green, n = 23). Bootstrap support values from 100 replicates are indicated at the nodes of the tree.

In the ADMIXTURE analysis, the lowest cross-validation errors [2] were always found for K = 3, indicating contribution of three ancestral populations (Figure 7). Majority of samples from each of the populations show a single dominant component of ancestry with little contribution from other populations, with the exception of four individuals from Haki, which show clear admixture of the Biaowiea population.

**Figure 7:**
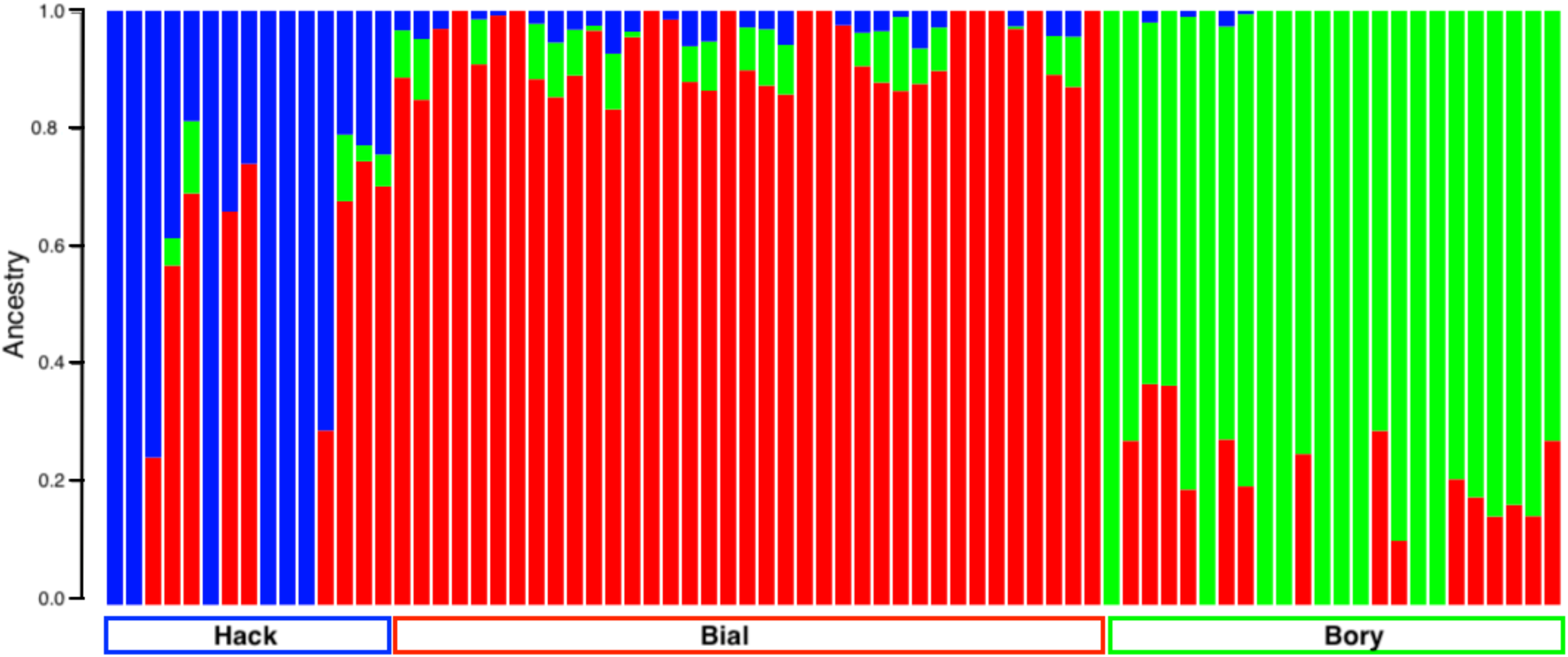
Maximum likelihood Admixture analysis of all *A. flavicollis* samples for the optimal K = 3. Each bar represents an individual and each colour represents its ancestry component (red: Biaowiea, blue: Haki, green: Bory Tucholskie).

Recognising that STRUCTURE-type analyses (on which ADMIXTURE is based) may be sensitive to the effects of uneven number of samples in compared groups [53], we repeated the ADMIXTURE analysis 10 times, each time randomly drawing the same number of individuals (n = 15) from each population. In all cases, the lowest cross-validation errors were found for K = 2, followed by K = 3 (Supplementary Figure S5). At even sampling, ADMIXTURE pattern found for K = 3 was the closest to the observed ecological and geographical distribution of the samples and closely matched our results when all samples were included (Supplementary Figure S6).

The patterns of heterozygosity highlight Haki as the only population where the values of H_o_ is higher than H_e_, where the F_IS_ is negative (Table 2). As parameters such as number of private alleles, nucleotide diversity and heterozygosity can vary with sample size, we performed 100 calculations of the above parameters using random sampling of the same number of individuals (n = 15) from each population. The parameters showed similar relationships except for the number of private alleles (data not shown).

**Table 2:** Population genetic parameters calculated based on 12654 SNPs from all 72 individuals of *A. flavicollis*. N, number of individuals; Npa, number of private alleles; Ind per loci, Mean number of individuals per locus in this population; H_o_ observed and H_e_ expected heterozygosity;, average nucleotide diversity; F_JS_ inbreeding coefficient

| Pop ID | N | Npa | Ind per loci | Obs Het | Exp Het | Pi | Fis |
| --- | --- | --- | --- | --- | --- | --- | --- |
| **Haki** | 15 | 32 | 14.42 | 0.30 | 0.27 | 0.28 | -0.04 |
| **Bory Tucholskie** | 24 | 74 | 22.93 | 0.28 | 0.28 | 0.29 | 0.02 |
| **Biaowiea** | 37 | 148 | 35.13 | 0.29 | 0.30 | 0.30 | 0.01 |

F_st_ values are consistently very low between all the populations, even though populations from Haki and Bory Tucholskie show three-fold higher F_st_ values that for the other two pairs of populations (Table 3).

**Table 3:** Pairwise F_ST_ values for the three populations of *A. flavicollis*.

|  | Bory Tucholskie | Biaowiea |
| --- | --- | --- |
| **Haki** | 0.085 | 0.055 |
| **Bory Tucholskie** |  | 0.045 |

### Species divergence

Finally, we calculated that the average p-distance between *A. flavicollis* and *A. sylvaticus*, based on 21377 shared loci, is 1.51 % (standard deviation = 1.11%).

We then identified the top 117 most divergent loci between the species, which all had the divergence larger than 4.9% (The loci ID are provided in the Supplementary Table S3), and checked whether these loci alone allow for accurate assignment of samples to the two species. We constructed PCA plots from the Polish samples only and from the Polish, other European and Tunisian samples together. They demonstrate that while the 117 loci are sufficient to clearly assign Polish samples to the two species (Supplementary Figure S8), some uncertainty remains when we use these loci for the broader set of samples. Whereas all *A. flavicollis* samples do cluster together, *A. sylvaticus* samples do not form a clearly differentiated group (Supplementary Figure S9).

We also identified fixed loci, where all individuals within each species have identical sequences. There were 3526 such fixed loci for *A. flavicollis* and 5843 for *A. sylvaticus*. We then used 1273 of those loci that were shared among the two species and calculated that the average p-distance based on fixed differences is 0.97% (standard deviation = 0.94%).

## Discussion

RAD-sequencing approaches, including double-digest RAD-seq and its variants [6, 18, 41, 48, 49], have allowed a cost-effective discovery of thousands of genetic markers in both model and non-model organisms [20, 59], proving to be a transformative research tool in population genetics [8, 13, 23], phylogeography and phylogenetics [4, 22, 26, 56], marker development [47], linkage mapping studies [7], species differentiation [44] and detecting selection [61]. However, despite the widespread use of this approach to marker discovery, only few studies have used RAD-seq in mammals [29, 31, 43, 60?]. Here, we have identified over 10000 markers in two closely related and common species of *Apodemus* in Western Palearctic, characterised the population structure of *A. flavicollis* and compared it to *A. sylvaticus*, for the first time providing estimates of the species divergence and population genetic parameters based on thousands of SNPs.

### Technical considerations

We have used four pairs of technical duplicates to check the accuracy of the RAD-seq genotyping based on the Poland protocol [50]. The largest source of discrepancy in SNP calls between the duplicates is caused by unequal identification of loci: the difference in our case averaged approximately 10% (Table 1) and was similar to allele misindentification rates. However, when considering only shared loci between the duplicates, the discrepancy in SNP calls fell by over an order of magnitude to an average of 0.5%, indicating high accuracy and reliability of calls in once-defined shared loci. Our finding of loci calls being the major source of genotyping variability agrees with Mastretta *et al*. (2015), although our discrepancies are almost an order of magnitude smaller. Moreover, despite the differences in number of loci included in the analysis, each duplicated pair of samples clustered together with a 100% bootstrap values support and branch length equal to 0 on the phylogenetic tree (Figure 6), indicating that the samples were identical. Overall, our finding reiterates the importance of the influence of stochastic events and imprecise size selection in the library preparations on genotyping calls [36]. We note that some of these variables could be better controlled with automated size-selection approaches [48]. Our findings also illustrate the usefulness of including technical replicates during library preparation.

### Effect of group size

Permutations performed for the calculations of genetic diversity parameters (Table 4) have shown that with the exception of the number of private alleles, the results are comparable, regardless of the number of samples included per each population.

**Table 4:** Average genetic diversity parameters for *Apodemus flavicollis* calculated from 100 permutations of 45 individuals (15 samples per population, 12654 SNPs). N, number of individuals; Npa, number of private alleles; Ind per loci, Mean number of individuals per locus in this population; H_o_ observed and H_e_ expected heterozygosity;, average nucleotide diversity; F_IS_ inbreeding coefficient

| Pop ID | N | Npa | Ind per loci | Obs Het | Exp Het | Pi | Fis |
| --- | --- | --- | --- | --- | --- | --- | --- |
| **Haky** | 15 | 115.53 | 14.42 | 0.31 | 0.28 | 0.29 | -0.05 |
| **Bory Tucholskie** | 15 | 183.95 | 14.33 | 0.29 | 0.28 | 0.29 | 0.02 |
| **Biaowiea** | 15 | 204.84 | 14.23 | 0.30 | 0.29 | 0.31 | 0.02 |

In the ADMIXTURE, we observed different optimal K depending on whether all samples were included in the analysis (K = 3) or a set of 15 randomly-chosen set of samples from each population (K = 2, although closely followed by K = 3). While the previously reported tendency of STRUCTURE-like analyses to produce ΔK = 2 does not apply in our case due to different method to select optimal number of clusters [25], we chose to use K = 3 for our analyses due to close match to the spatial and ecological locations from which our populations were sampled. The results obtained for K = 3 in the evenly-sampled dataset were similar to the clusters obtained for K = 3 with the complete dataset. ADMIXTURE plots for K = 2 to K = 5 are shown as Supplementary Figure S7.

### Population structure

The F_ST_ values calculated in this study between all three pairs of populations of *A. flavicollis*, based on 12654 SNPs, are consistently low and are not affected when we randomly draw the same number of individuals from each population to compute pairwise F_ST_ (Table 5). Previous studies of *A. flavicollis* populations in north-eastern Poland based on a small number of microsatellites showed similarly and consistently low values [15, 19], even though Gortat et al. [19] suggested some population structure based on statistically significant differences between very low pairwise F_ST_ values. Czarnomska et al. [15] also suggest large, broadly geographically defined clusters of *A. flavicollis* in north-eastern Poland that are separated by highly admixed individuals, but, again, F_ST_ between those clusters are as low as those reported by Gortat et al. [19] and this study.

**Table 5:** Average pairwise F_ST_ values ś standard deviation for the three populations of *A. flavicollis* calculated from 100 permutations of 45 individuals (15 samples per population, 12654 SNPs)

|  | Bory Tucholskie | Biaowiea |
| --- | --- | --- |
| **Haky** | 0.086 ± 0.002 | 0.057 ± 0.001 |
| **Bory Tucholskie** | | 0.045 ± 0.002 |

We would argue, based on a much larger set of markers reported here, that *A. flavicollis* has a negligible population structure across the entire area studied. Large number of markers nevertheless allows us to discover evidence for admixture of Biaowiea population and Haki (Figure 7), further indicated by relatively high heterozygosity and negative F_IS_ in this population. It is therefore intriguing that such a low differentiation occurs across hundreds of kilometres of varying landscape in a species that typically has a limited range of about 4 km^2^ and that suffers up to 86% winter mortality rate [52], which would lead to multiple bottlenecks and drift-driven population differentiation. With this in mind, our data suggests a much larger dispersal ability of the species, a much better connectivity between populations, or both.

Both low overall F_ST_ and moderate heterozygosity suggest it would be worthwhile to conduct a genome-wide scan for selection using F_ST_ as a metrics of local genomic differentiation to identify geographically local regions under selection. This, however, is not yet possible given the lack of high-quality reference genome for *Apodemus* and unknown synteny to the available genome of *Mus musculus*.

### Divergence and differentiation of *A. flavicollis* and *A. sylvaticus*

Given that accurate identification of the two species using morphological characters is problematic, especially in their southern range [10], a large collection of markers identified in this study allowed us to create a catalogue of 632060 loci that allow clear differentiation between species. This identification is somewhat biased, as the catalogue was built using many more samples of *A. flavicollis* than *A. sylvaticus* (72 vs 10) and both from a relatively limited geographical range. Nevertheless, it allowed for accurate assignment of *A. flavicollis* samples and to a large degree of *A. sylvaticus*, as we demonstrated on a set of 20 independent samples from other European countries and Tunisia (Figure 4). The imperfect clustering of some European *A. sylvaticus* samples on Figure 4 may reflect the higher genetic differentiation of population from the edge of the range of the species: samples near the centre of the PC1 axis on Figure 4 (green dots in a circle) all come from Tunisia. Given the wide distribution of both species in Western Palearctic, a more representative sample from both species from a broader geographic range would likely provide more accurate set of markers for their identification.

Finally, we calculated the nucleotide divergence between the two species, based on 21377 shared loci, which is 1.51%. Considering a divergence time between *A. flavicollis* and *A. sylvaticus* estimated from archeological data of 4 Mya [37], the evolutionary rate is 0.0019 substitutions per site per million of years. While this estimate of sequence divergence level is in broad agreement with calculations based on mitochondrial 12S rRNA, IRBP and Cytochrome b genes [38], we note that Aghova *et al*. (2018) [1] provided an updated timing of a split between *A. sylvaemus* and *A. mystacinus*, which, at 9.6Mya, is 2Mya older than previously suggested by Michaux *et al*. (2002) [38]. If we use that date as a reference and move the presumed split between *A. flavicollis* and *A. sylvaticus* by 2.6Mya, the estimated evolutionary rate from our data would be 0.0011. In both cases, our calculation is likely an under-estimate, as we only used shared loci to calculate divergence and did not include the potential impact of insertion/deletion events, which can significantly affect the total genomic divergence between species [9, 34]. Highly divergent sequences would have been identified as different loci, and would not be compared to their true homologous sequences.

## Conclusions

We have successfully applied the ddRAD-seq approach to discover tens of thousands of SNPs in wide-spread and common mammalian species of *A. flavicollis* and *A. sylvaticus*. The high resolution data obtained here allowed us to delineate geographically close populations, including identifying admixture between them, but suggest that *A. flavicollis* effectively forms a single population in an entire sampling area that spans 500 km in the W-E direction. Comparing *A. flavicollis* and *A. sylvaticus*, we have calculated their divergence and identified a set of genomic loci that enable effective molecular identification of the species. We anticipate that with the development of further whole-genome resources, *Apodemus*, thanks to its common status, broad geographic range and long history of ecological observations, will become an excellent model species for evolutionary and ecological research in the genomic era.

## Methods

### Sample collection and DNA extraction

Eighty two individuals (10 *Apodemus sylvaticus* and 72 *Apodemus flavicollis*) from four locations in northern Poland spanning 500 km were trapped in 2015 (Figure 8). *A. flavicollis* were collected in Biaowiea (E23.8345814, N52.7231935), an oak-lime-hornbeam forest (n = 35), Bory Tucholskie (E17.5160265, N53.7797608), in an oak-lime-hornbeam and pine forest (n = 23) and Haki (E23.1793284, N52.834369), in a xerothermic meadow (n = 14). *A. sylvaticus* were trapped in Kadzido (E21.3778496, N53.2089113) in a dry pine forest (n = 5) and in Bory Tucholskie, mainly in a pine forest (n = 5) (Supplementary Table S1). While *A. flavicollis* are present in all sampled locations, there have been no trappings of *A. sylvaticus* in Biaowiea for the last 20 years, despite Biaowiea being within the European range of this species (Dr Karol Zub, personal communication). The sampling procedures were approved by the Local Ethical Commission on Experimentation on Animals in Biaystok, Poland, under permission number 2015/99. All animals were released after sample collection.

**Figure 8:**
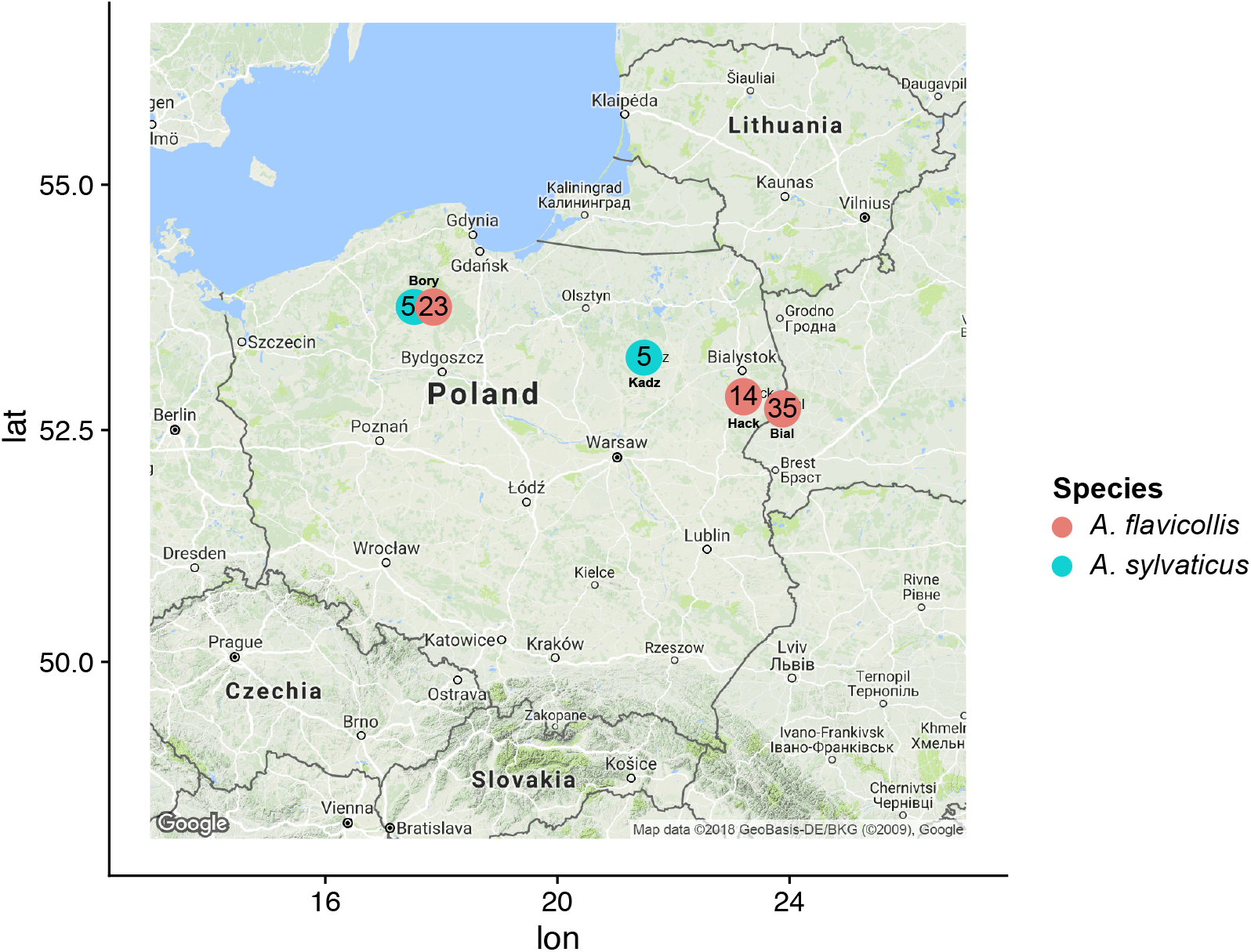
Locations of the Polish samples used in this study. Red circles represent samples from *Apodemus flavicollis* while blue dots represent samples from *Apodemus sylvaticus*. The number inside the circles are the number of samples from each locality. Bial - Biaowiea, Kadz - Kadzido, Hack - Haki, Bory - Bory Tucholskie.

Tail clippings were collected, preserved in ≥ 95% ethanol and stored at −20°C until DNA extraction. The tissues were digested by incubating at 55°C overnight with lysis buffer (10mM Tris, 100mM NaCl, 10mM EDTA, 0.5% SDS) and proteinase K (20mg/ml). Subsequently, potassium acetate and RNAse A were used to remove protein and RNA contamination. Three ethanol washes were performed using Sera-Mag SpeedBeads solution (GElifesciences, Marlborough, MA, USA). The quality and integrity of the DNA was tested in a 2% agarose gel. Twenty-fold dilutions of the samples were used to measure the DNA concentration using Quant-iT PicoGreen dsDNA assay kit (Invitrogen, Carlsbad, CA, USA) and concentration of each sample was then normalised to 10 ng/*μ*l in 20*μ*l volume. Four samples were used as technical duplicates (F06-B02, G02-D01, H11-G06, F12-A12). Technical duplicates had the same DNA but were digested and ligated to barcodes independently.

### ddRAD-seq library preparation

ddRAD-seq library was prepared following the protocol from Poland and Rife [49], adapted to a different combination of enzymes. Briefly, genomic DNA was digested in a 20 *μl* reaction with CutSmart^®^ buffer, 8 units of *SbfI* and 8 units of HF-*MseI* (New England Biolabs, Frankfurt am Main, Germany). Digestion was performed at 37°C for 2 hours. Enzymes were inactivated at 65°C for 20 minutes and the reactions were kept at 8°C. Adapter ligation was performed at 22°C for 2 hours and the ligase was inactivated by incubating the samples at 65°C for 20 minutes. Samples were cooled down to 8°C and multiplexed by combining 5*μl* of each sample. P1 adapters contained barcodes with a length between 5 and 10 bp.

PCR amplification was conducted in 25*μl* with 1*μl* of each primer (IlluminaF_PE: AATGATACGGCGACCACCGAGATCTACACTCTTTCCCTACAC-GACGCTCTTCCGATCT and IlluminaR_PE: CAAGCAGAAGACGGCATAC-GAGATCGGTCTCGGCATTCCTGCTGAA) at 10mM, 0.5*μl* of 10 mM dNTPs, 13.25*μl* of PCR-grade water, 5*μl* of 5x Phusion HF Buffer, 0.25*μl* of Phusion DNA Polymerase (New England Biolabs, Frankfurt am Main, Germany) and 4*μl* of the multiplexed DNA. After an initial denaturation step of 30s at 98°C, PCR reaction was carried out for 12 cycles (10s at 98°C, 20s at 58°C and 15s at 72°C). Final elongation step was performed at 72°C for 5 minutes.

PCR products were loaded into a single lane on a 1% agarose gel with 100 bp DNA ladder (New England Biolabs, Frankfurt am Main, Germany). Fragments between 200 and 500 bp were cut from the gel with a scalpel and purified using the QIAquick gel extraction kit (QIAgen, Hilden, Germany), followed by the second cleanup step with Sera-Mag SpeedBeads (GElifesciences, Marlborough, MA. USA). Sizing, quantification and quality control of the DNA was performed using Bioanalyzer 2100 (Agilent, Santa Clara, CA, USA) before paired-end sequencing on an Illumina HiSeq 3500 with 150 bp read length.

### Processing of RAD-tags

Sequences were analysed with Stacks version 1.48 [11]. Samples were demultiplexed using process_radtags allowing no mismatches in barcodes and cutting sites. Sequences with uncalled bases and low quality scores were removed and all reads were trimmed to 141 bp. The four files generated per sample by process_radtags were concatenated using a custom bash script. The best parameters for building and calling SNPs *de novo*, using denovo_map, were calculated following Paris et al. [46] approach, using either samples from both species or only from *A. flavicollis*. Secondary reads were not used to call haplotypes in denovo_map (option -H).

### SNPs and loci co-identification rates

We estimated the loci and SNP co-identification rates by analysing a set of four samples that were prepared and sequenced in duplicates. Sequences for 52494 loci from both species, were extracted using –fasta_samples option from the population package in Stacks. We extracted sequences for each of the duplicated samples with a custom script and calculated co-identification rates as described by [36]. Briefly, the locus misassignment rate is the percentage of unidentified loci, calculated by dividing the number of loci found only in one of the duplicates by the total number of loci in each sample. The allele misassignment rate is the percentage of mismmatches between the IUPAC consensus sequences between homologous loci from each pair of duplicates. Finally, the two SNP error rates: the percentage of different SNPs called in each of the duplicated samples using either all 10178 SNPs or using the SNPs called without missing data between duplicate samples excluded (see Table 1).

### Variant calling and filtering

We combined the data from *A. sylvaticus* and *A. flavicollis* to establish species differentiation and then filtered the SNPs using the population package from Stacks [11] and VCFtools [16]. We kept SNPs from the loci present in the 80% of the individuals in each species (p=1, r=0.8) and excluded SNPs with minor allele frequencies MAF<0.05 and which deviated from the Hardy-Weinberg equilibrium (HWE) at P<0.05. We also removed sites with mean depth values lower than 20. We manually modified the chromosome numbers in the vcf file to input it into SNPhylo [33], which we used to build the tree. We set a missing rate (-M) of 1, minor allele frequencies (-m) of 0, linkage disequilibrium threshold (-l) of 1 and the -r option to skip the step of removing low quality data. Confidence values were estimated using 1000 bootstrap replicates. The root was manually fixed to separate both species. Principal Component Analysis (PCA) was performed using the R package Adegenet [28] (Figure 3).

The set of divergent loci identified between the two species in Polish samples was tested for its ability to differentiate an extra set of samples from other locations in Europe and Tunisia. Ten *A. flavicollis* (2 samples from Austria, 5 from Lithuania and 3 from Romania) and 10 samples of *A. sylvaticus* (4 samples from Wales, 3 from Tunisia and 3 from Scotland) were kindly provided by Dr Jeremy Herman, National Museums Scotland, Dr Johan Michaux, University of Liege and Dr Karol Zub, Mammal Research Institute of the Polish Academy of Sciences (MRI) (Supplementary Table S4). We considered all 20 test samples as a different group from Polish *A. sylvaticus* and *A. flavicollis* for SNP calling. We kept SNPs from the loci present in the 80% of the individuals in each group (p=1, r=0.8) and excluded SNPs with minor allele frequencies MAF<0.05, SNPs which deviated from the Hardy-Weinberg equilibrium (HWE) at P<0.05, sites with mean depth values lower than 20 and with more than 5% of missing data.

### Population divergence

To analyse genetic diversity and population connectivity within *A. flavicollis*, we analysed the three populations (Bory Tucholskie, Biaowiea and Haki) separately (p=3, r=0.8), while keeping the other parameters as described above. We note that absence of filtering out SNPs in Hardy-Weinberg disequlibrium did not markedly affect the reported pairwise F_st_ values (data not shown). Due to the lack of outgroup, a mid-point root was chosen in the phylogenetic tree. Individual ancestries were estimated following a maximum likelihood approach with ADMIXTURE [3], after conversion of the VCF file to ped with plink version 1.9 [12, 54]. ADMIXTURE analysis was run for each of K=1 to K=5, each using 10 different seeds. Weighted (Weir-Cockerham) F_st_ was calculated with VCFtools v0.1.13. Heterozygosity, Pi and Fis were calculated with the population package from Stacks [11].

### Species divergence

To calculate the p-distance between the two species, first a set of common loci was extracted with a custom script and the strict consensus sequences for each species were calculated with Consensus.pl script [24] with no threshold parameter set, such that the most common nucleotide was set as a consensus. p-distance was then calculated using a custom R script that counts the number of differences between pairs of sequences (one from each species, for each of the 21377 loci) (Supplementary Materials, Section 9). The same set of scripts was also used to select loci with fixed differences within each species and then to calculate the p-distance between them.

## Supporting information

biorxiv_supp_files.pdf

## Abbreviations

mtDNA: mitochondrial DNA
ddRAD-seq: double-digest restriction site-associated DNA sequencing
SNP: single nucleotide polymorphism
MAF: minor allele frequency
HWE: Hardy-Weinberg equilibrium
PCA: principal component analysis
Fst: fixation index
VCF: variant call format

## Declarations

### Ethics approval and consent to participate

The sampling procedures were approved by the Local Ethical Commission on Experimentation on Animals in Biaystok, Poland, under permission number 2015/99.

### Consent for publication

Not Applicable.

### Availability of data and materials

The sequence data, in the format of the output from the Stacks 1.48 *proces_radtags* (four files per sample), have been deposited in EBI SRA repository under accession number PRJNA554851. The scripts used in the analysis are deposited at GitHub.com, with links available in the Supplementary Materials.

### Author’s contributions

MLMC designed the study, prepared the sequencing library, analysed the data and wrote the manuscript. MK contributed to the experimental design and preparation of the sequencing library. KZ contributed to the experimental design, performed the sampling and contributed to the manuscript. YFC contributed to the analysis of the data and the manuscript. JB designed the study, analysed the data and wrote the manuscript.

## Acknowledgements

The authors wish to thank Jeremy Herman (National Museums Scotland) and Johan Michaux (Université de Liège) for providing the European/Tunesian samples for testing our catalog of SNPs. MLMC and JB also wish to acknowledge the use of the Orion High Performance Computing cluster at the School of Applied Sciences, University of Huddersfield.

## Additional Files

Additional file 1 — Supplementary Materials

**Supplementary Figure 1.**
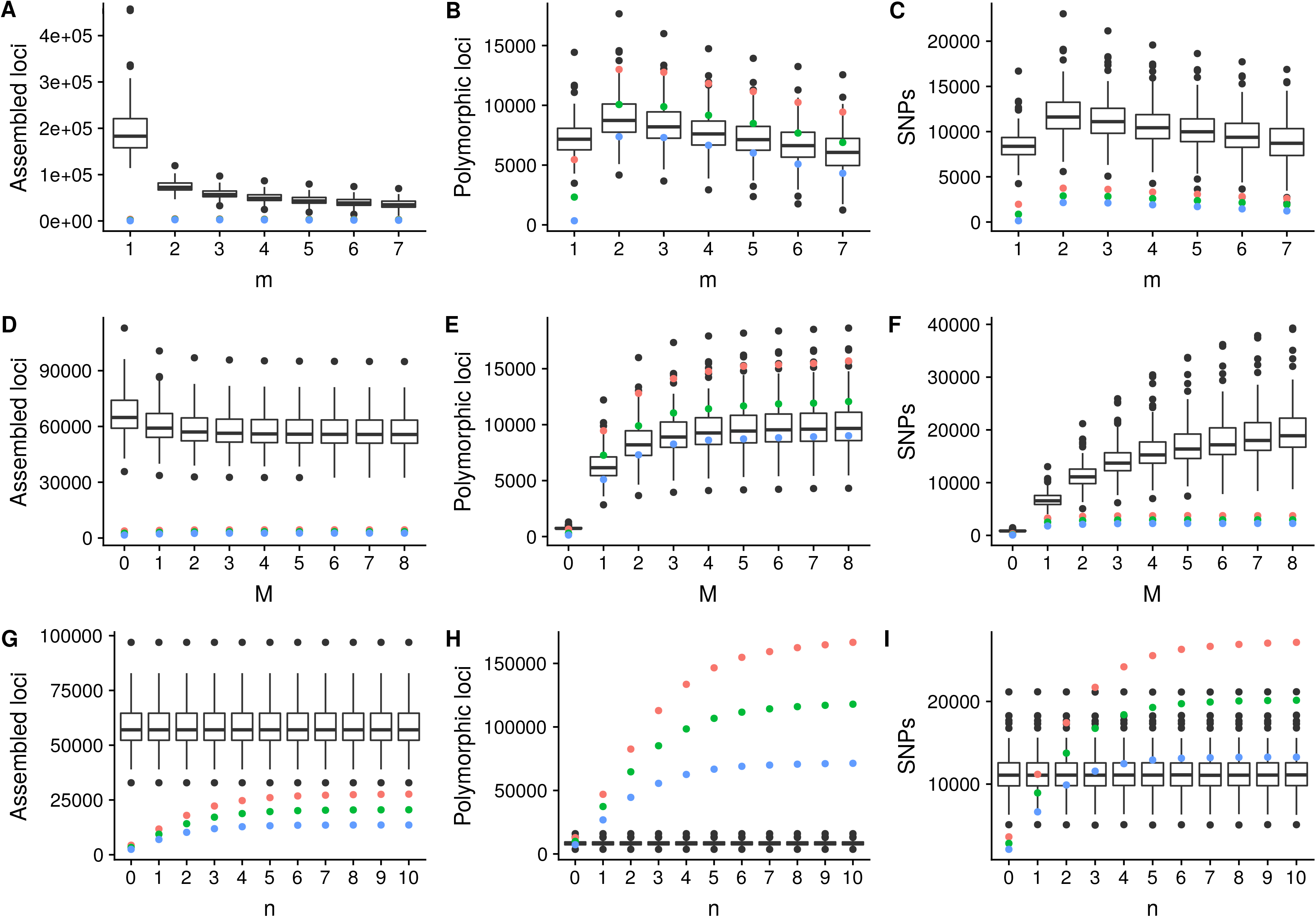

**Supplementary Figure 2.**
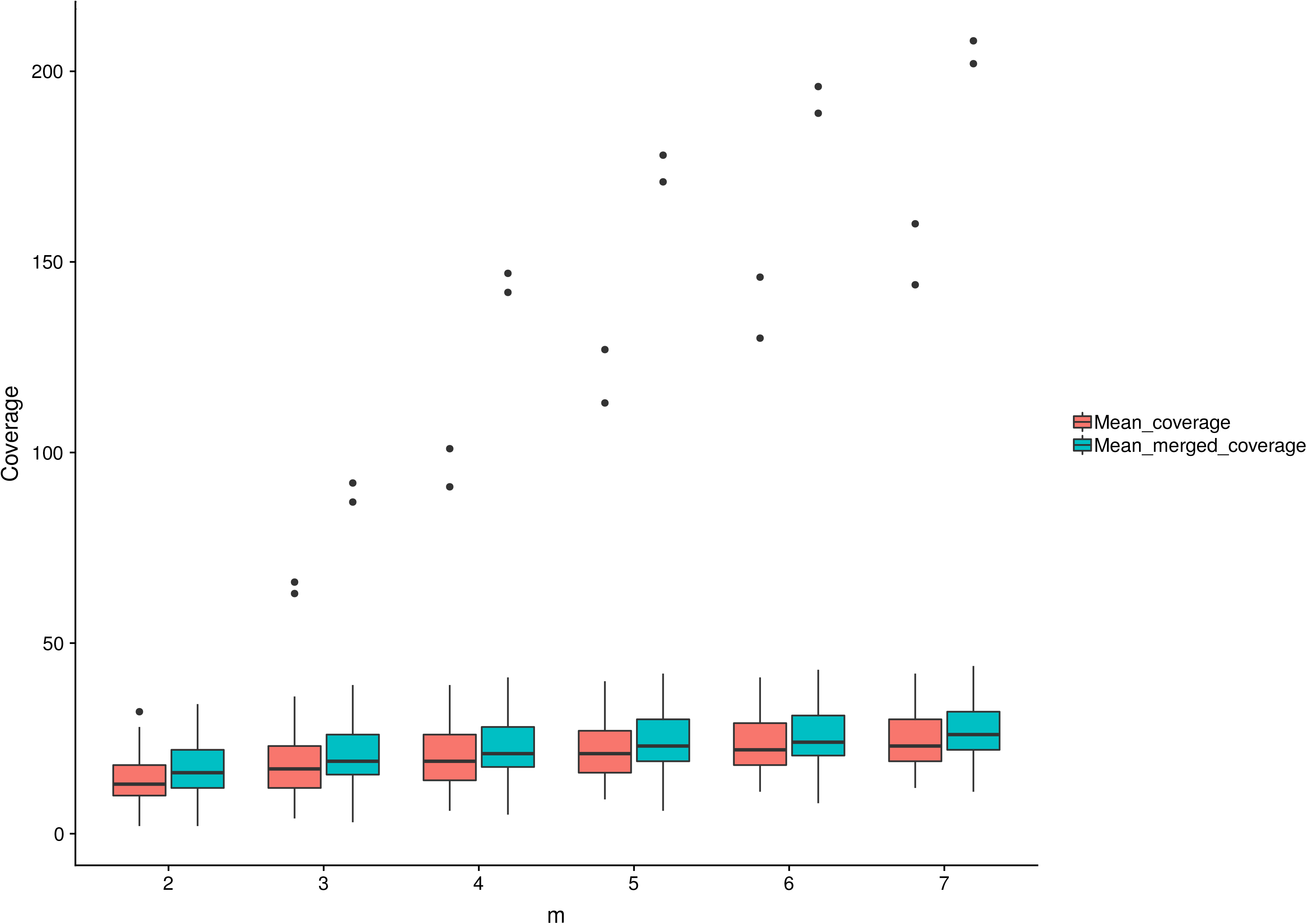

**Supplementary Figure 3.**
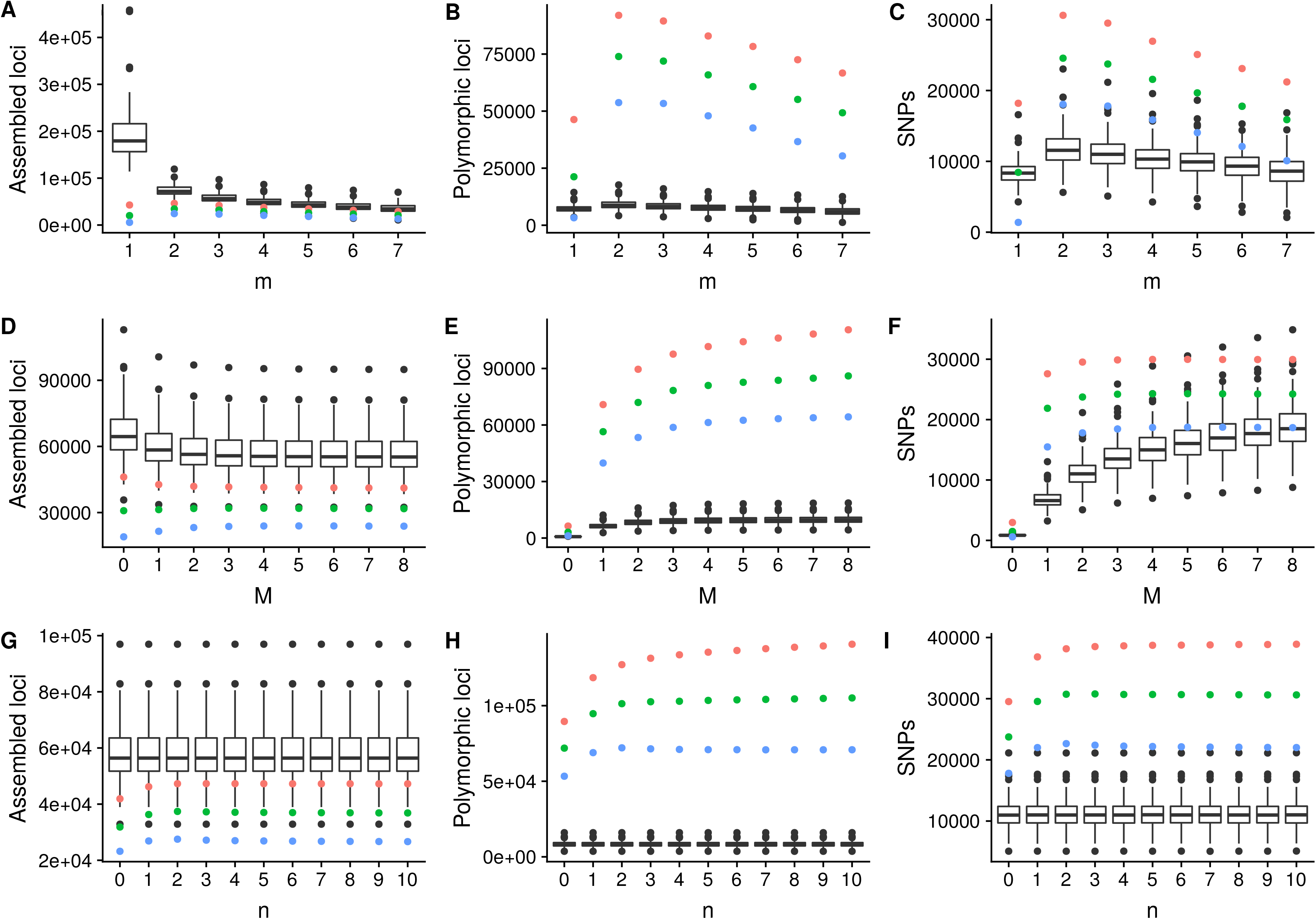

**Supplementary Figure 4.**
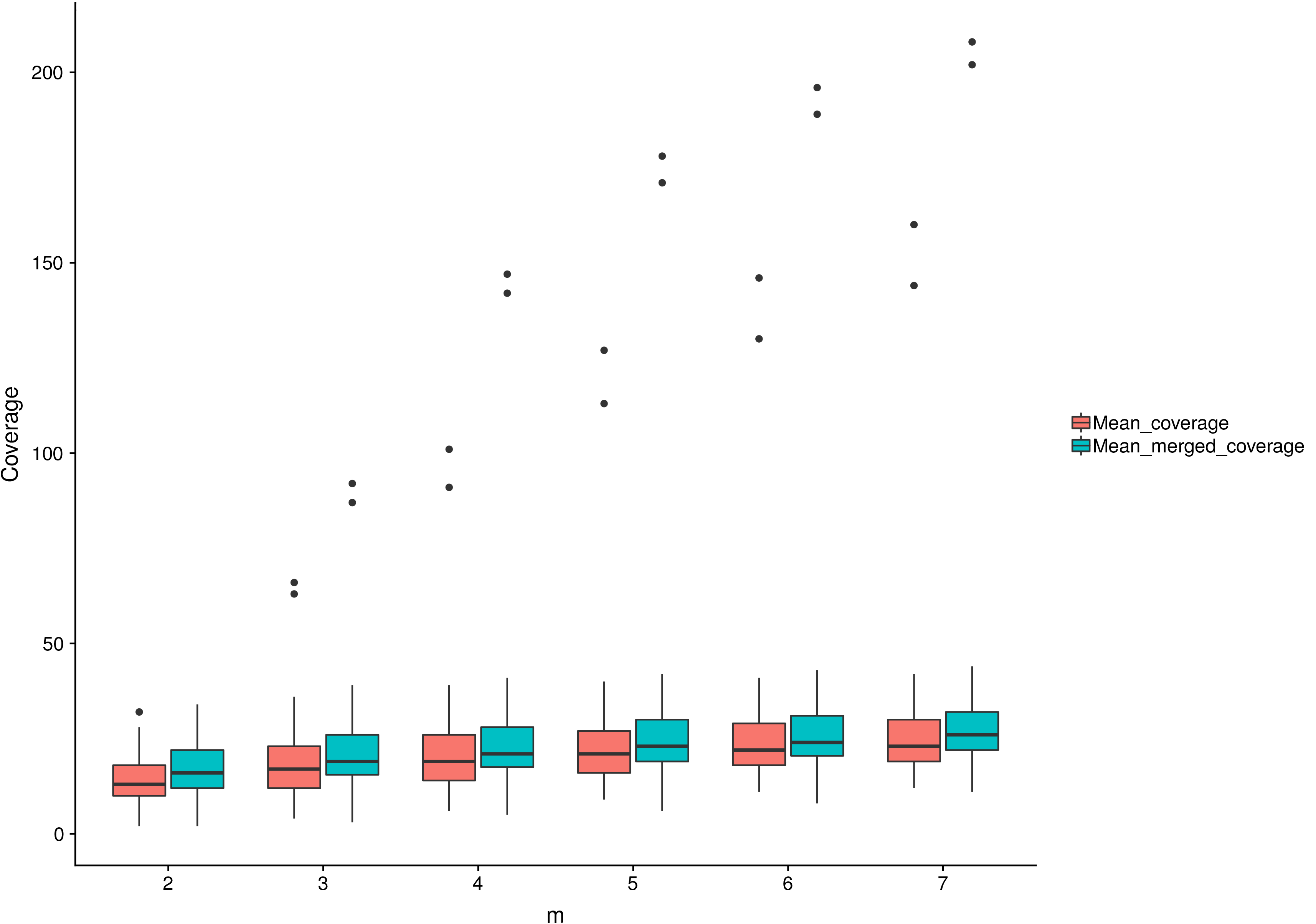

**Supplementary Figure 5.**
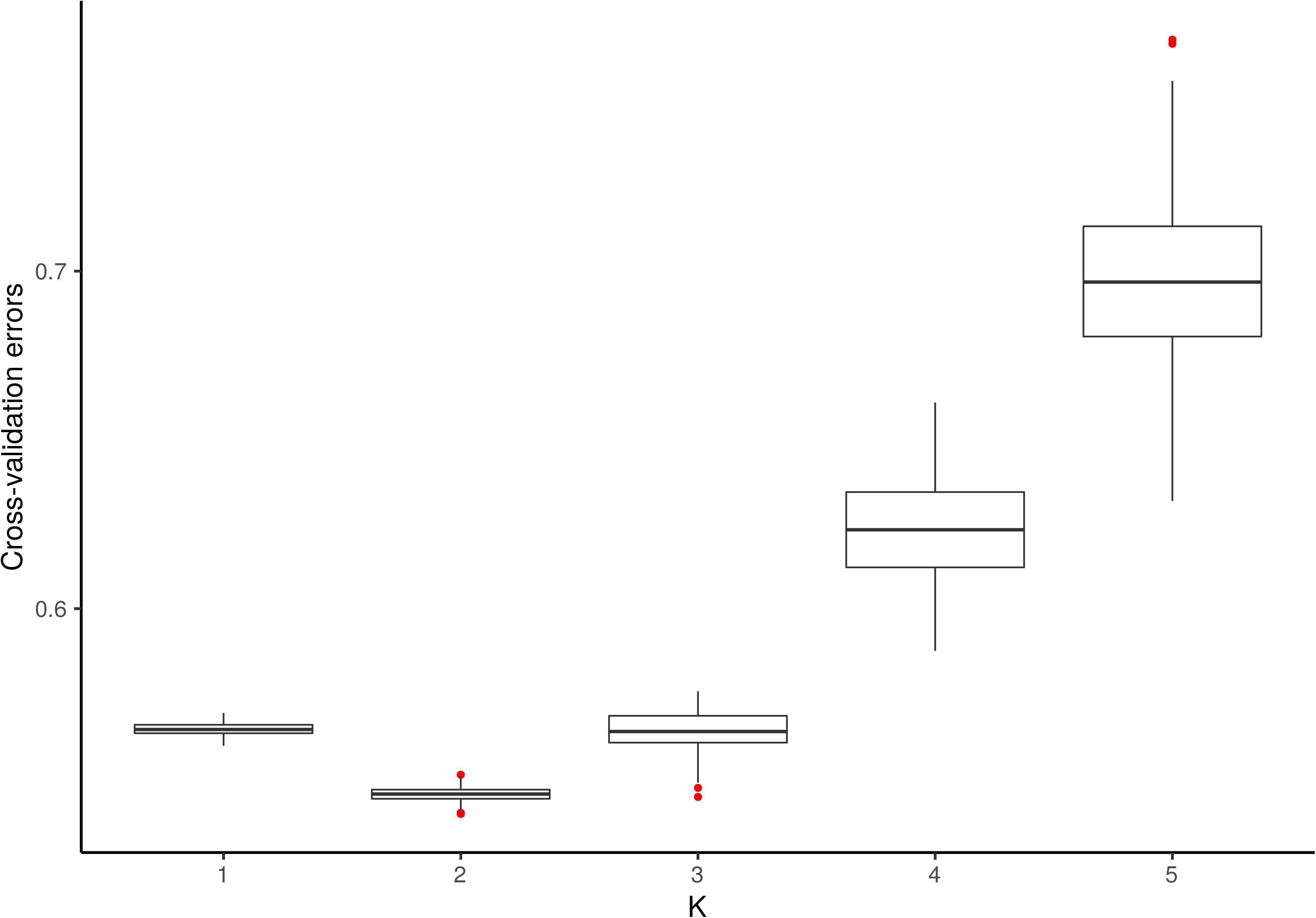

**Supplementary Figure 6.**
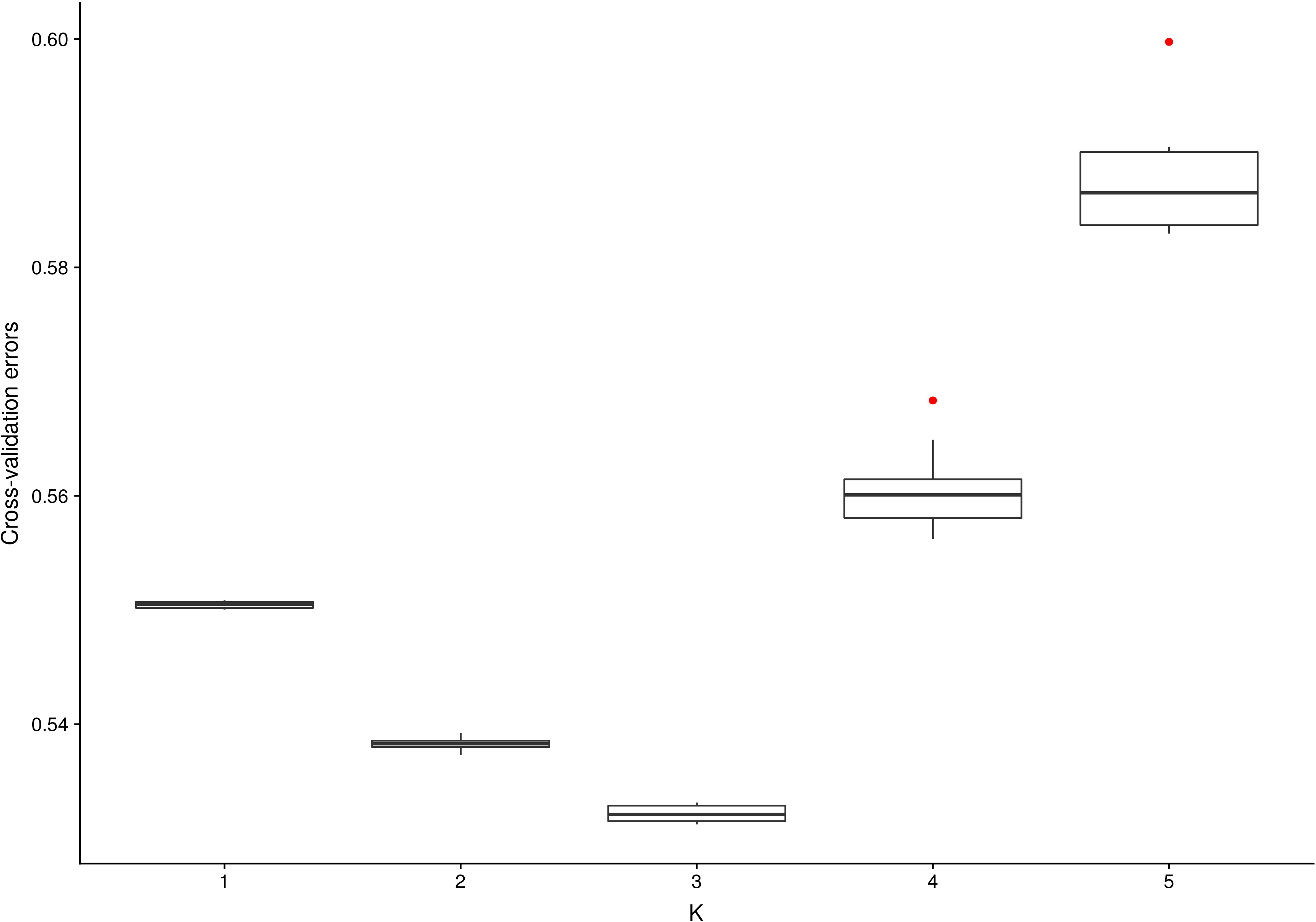

**Supplementary Figure 7.**
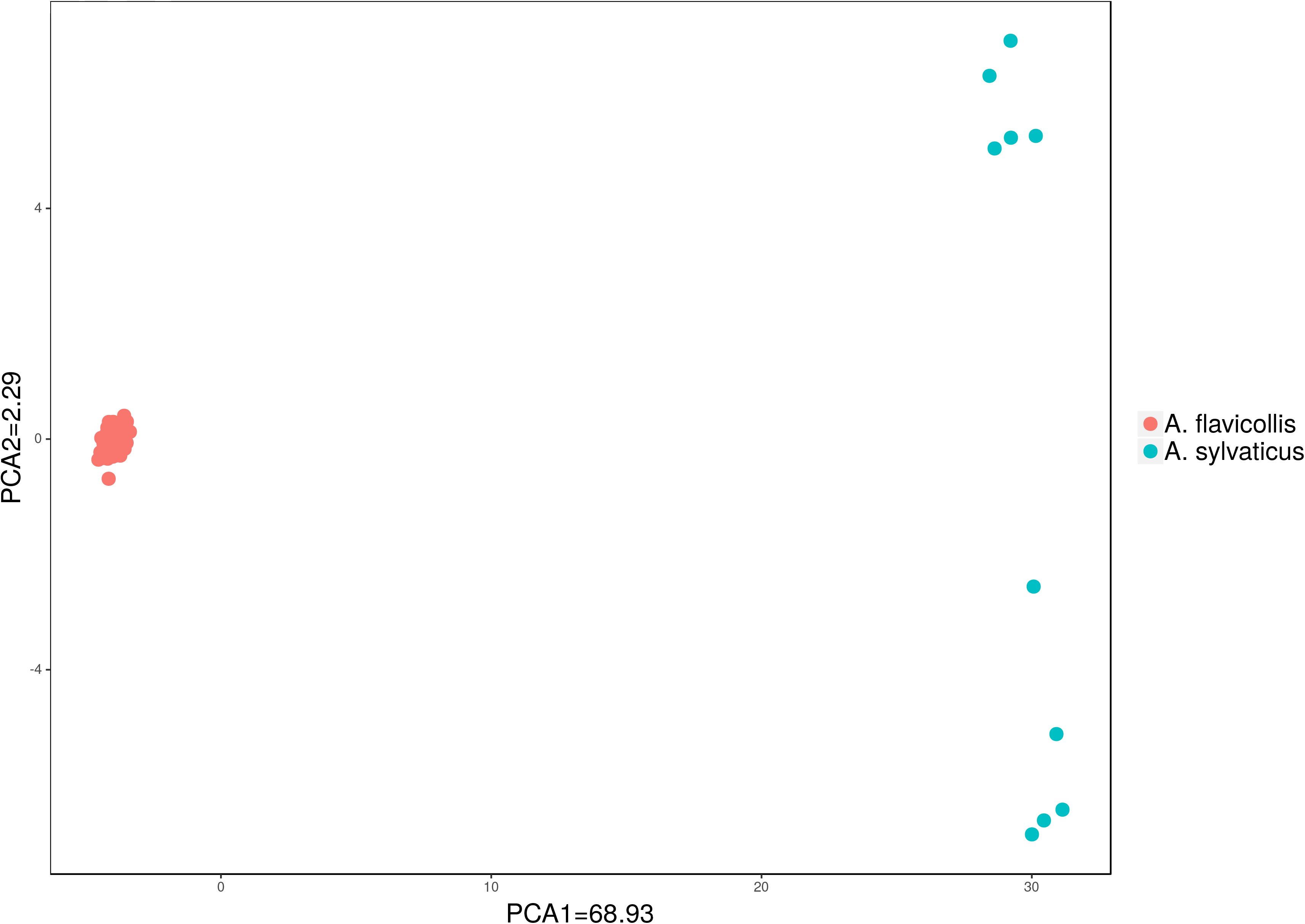

**Supplementary Figure 8.**
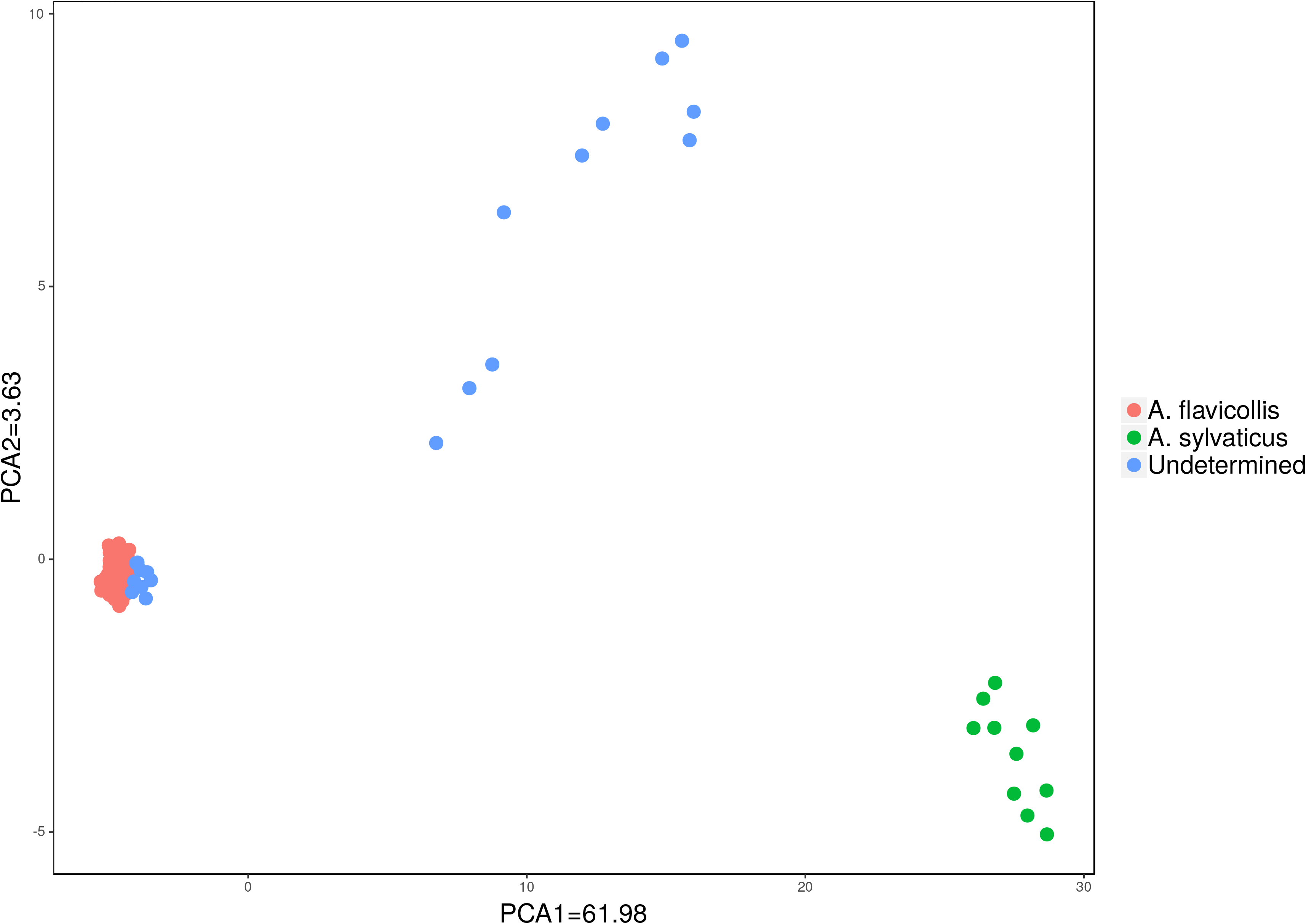

## References

1. Tatiana Aghová, Yuri Kimura, Josef Bryja, Gauthier Dobigny, Laurent Granjon, and Gael J. Kergoat. Fossils know it best: Using a new set of fossil calibrations to improve the temporal phylogenetic framework of murid rodents (Rodentia: Muridae). Molecular Phylogenetics and Evolution, 128:98–111, 2018.

2. David H Alexander and Kenneth Lange. Enhancements to the admixture algorithm for individual ancestry estimation. BMC bioinformatics, 12(1):246, 2011.

3. David H Alexander, John Novembre, and Kenneth Lange. Fast model-based estimation of ancestry in unrelated individuals. Genome research, 19(9):1655–1664, 2009.

4. S Elizabeth Alter, Jason Munshi-South, and Melanie LJ Stiassny. Genome wide snp data reveal cryptic phylogeographic structure and microallopatric divergence in a rapids-adapted clade of cichlids from the congo river. Molecular ecology, 26(5):1401–1419, 2017.

5. Kimberly R Andrews, Jeffrey M Good, Michael R Miller, Gordon Luikart, and Paul A Hohenlohe. Harnessing the power of radseq for ecological and evolutionary genomics. Nature Reviews Genetics, 17(2):81, 2016.

6. N.A. Baird, P.D. Etter, T.S. Atwood, M.C. Currey, A.L. Shiver, Z.A. Lewis, E.U. Selker, W.A. Cresko, and E.A. Johnson. Rapid snp discovery and genetic mapping using sequenced rad markers. PloS one, 3(10):e3376, 2008.

7. Simon W Baxter, John W Davey, J Spencer Johnston, Anthony M Shelton, David G Heckel, Chris D Jiggins, and Mark L Blaxter. Linkage mapping and comparative genomics using next-generation rad sequencing of a non-model organism. PloS one, 6(4):e19315, 2011.

8. L Blanco-Bercial and A Bucklin. New view of population genetics of zooplankton: Rad-seq analysis reveals population structure of the north atlantic planktonic copepod centropages typicus. Molecular ecology, 25(7): 1566–1580, 2016.

9. Roy J Britten. Divergence between samples of chimpanzee and human dna sequences is 5%, counting indels. Proceedings of the National Academy of Sciences, 99(21):13633–13635, 2002.

10. Vanja Bugarski-Stanojević, Jelena Blagojević, Tanja Adnadjević, Vladimir Jovanović, and Mladen Vujošević. Identification of the sibling species apodemus sylvaticus and apodemus flavicollis (rodentia, muridae)comparison of molecular methods. Zoologischer Anzeiger-A Journal of Comparative Zoology, 252(4):579–587, 2013.

11. Julian M Catchen, Angel Amores, Paul Hohenlohe, William Cresko, and John H Postlethwait. Stacks: building and genotyping loci de novo from short-read sequences. G3: Genes, genomes, genetics, 1(3):171–182, 2011.

12. Christopher C Chang, Carson C Chow, Laurent CAM Tellier, Shashaank Vattikuti, Shaun M Purcell, and James J Lee. Second-generation plink: rising to the challenge of larger and richer datasets. Gigascience, 4(1):7, 2015.

13. Gareth A Cromie, Katie E Hyma, Catherine L Ludlow, Cecilia Garmendia-Torres, Teresa L Gilbert, Patrick May, Angela A Huang, Aimée M Dudley, and Justin C Fay. Genomic sequence diversity and population structure of saccharomyces cerevisiae assessed by rad-seq. G3: Genes, Genomes, Genetics, 3(12):2163–2171, 2013.

14. Benjamin Cull, Alexander GC Vaux, Lisa J Ottowell, Emma L Gillingham, and Jolyon M Medlock. Tick infestation of small mammals in an english woodland. Journal of Vector Ecology, 42(1):74–83, 2017.

15. Sylwia D Czarnomska, Magdalena Niedziałkowska, Tomasz Borowik, and Bogumia Jędrzejewska. Regional and local patterns of genetic variation and structure in yellow-necked mice-the roles of geographic distance, population abundance, and winter severity. Ecology and evolution, 8:8171–8186, 2018.

16. Petr Danecek, Adam Auton, Goncalo Abecasis, Cornelis A Albers, Eric Banks, Mark A DePristo, Robert E Handsaker, Gerton Lunter, Gabor T Marth, Stephen T Sherry, et al. The variant call format and vcftools. Bioinformatics, 27(15):2156–2158, 2011.

17. Jamshid Darvish, Zeinolabedin Mohammadi, Fatemeh Ghorbani, Ahmad Mahmoudi, and Sylvain Dubey. Phylogenetic relationships of apodemus kaup, 1829 (rodentia: Muridae) species in the eastern mediterranean inferred from mitochondrial dna, with emphasis on iranian species. Journal of mammalian evolution, 22(4): 583–595, 2015.

18. Paolo Franchini, Daniel Monné Parera, Andreas F Kautt, and Axel Meyer. quaddrad: a new high-multiplexing and pcr duplicate removal ddrad protocol produces novel evolutionary insights in a nonradiating cichlid lineage. Molecular Ecology, 26(10):2783–2795, 2017.

19. Tomasz Gortat, Alicja Gryczyńska-Siemiątkowska, Robert Rutkowski, Anna Kozakiewicz, Antoni Mikoszewski, and Michał Kozakiewicz. Landscape pattern and genetic structure of a yellow-necked mouse apodemus flavicollis population in north-eastern poland. Acta Theriologica, 55(2):109–121, 2010.

20. Nicholas M Hammerman, Ramon E Rivera-Vicens, Matthew P Galaska, Ernesto Weil, Richard S Appledoorn, Monica Alfaro, and Nikolaos V Schizas. Population connectivity of the plating coral agaricia lamarcki from southwest puerto rico. Coral Reefs, 37(1):183–191, 2018.

21. Jeremy S Herman, Fríđa Jóhannesdóttir, Eleanor P Jones, Allan D McDevitt, Johan R Michaux, Thomas A White, Jan M Wójcik, and Jeremy B Searle. Post-glacial colonization of europe by the wood mouse, apodemus sylvaticus: evidence of a northern refugium and dispersal with humans. Biological Journal of the Linnean Society, 120(2):313–332, 2017.

22. Andrew L Hipp, Deren AR Eaton, Jeannine Cavender-Bares, Elisabeth Fitzek, Rick Nipper, and Paul S Manos. A framework phylogeny of the american oak clade based on sequenced rad data. PLoS One, 9(4):e93975, 2014.

23. Paul A Hohenlohe, Mitch D Day, Stephen J Amish, Michael R Miller, Nick Kamps-Hughes, Matthew C Boyer, Clint C Muhlfeld, Fred W Allendorf, Eric A Johnson, and Gordon Luikart. Genomic patterns of introgression in rainbow and westslope cutthroat trout illuminated by overlapping paired-end rad sequencing. Molecular ecology, 22(11):3002–3013, 2013.

24. Joseph Hughes. Sequence-manipulation. GitHub repository, 2011.

25. Jasmine K Janes, Joshua M Miller, Julian R Dupuis, René M Malenfant, Jamieson C Gorrell, Catherine I Cullingham, and Rose L Andrew. The k= 2 conundrum. Molecular Ecology, 26(14):3594–3602, 2017.

26. Daniel L Jeffries, Gordon H Copp, Lori Lawson Handley, K Håkan Olsén, Carl D Sayer, and Bernd Hänfling. Comparing radseq and microsatellites to infer complex phylogeographic patterns, an empirical perspective in the crucian carp, carassius carassius, l. Molecular ecology, 25(13):2997–3018, 2016.

27. Vida Jojić, Vanja Bugarski-Stanojević, Jelena Blagojević, and Mladen Vujošević. Discrimination of the sibling species apodemus flavicollis and a. sylvaticus (rodentia, muridae). Zoologischer Anzeiger-A Journal of Comparative Zoology, 253(4):261–269, 2014.

28. Thibaut Jombart. adegenet: a r package for the multivariate analysis of genetic markers. Bioinformatics, 24 (11):1403–1405, 2008.

29. L Lacey Knowles, Rob Massatti, Qixin He, Link E Olson, and Hayley C Lanier. Quantifying the similarity between genes and geography across alaska’s alpine small mammals. Journal of Biogeography, 43(7): 1464–1476, 2016.

30. Marcin Kolodziej, Alicja Melgies, Justyna Joniec-Wiechetek, Aleksander Michalski, Anna Nowakowska, Grzegorz Pitucha, and MarcinN Niemcewicz. First molecular characterization of dobrava-belgrade virus found in apodemus flavicollis in poland. Annals of Agricultural and Environmental Medicine, 25(2):368–373, 2018.

31. Hayley C Lanier, Rob Massatti, Qixin He, Link E Olson, and L Lacey Knowles. Colonization from divergent ancestors: glaciation signatures on contemporary patterns of genomic variation in collared pikas (ochotona collaris). Molecular Ecology, 24(14):3688–3705, 2015.

32. A.G. Lapinski, M.V. Pavlenko, L.L. Solovenchuk, and V.V. Gorbachev. Some limitations in the use of the mitochondrial dna cytb gene as a molecular marker for phylogenetic and population-genetic studies by the example of the apodemus genus. Russian Journal of Genetics: Applied Research, 6(1):84–90, 2016.

33. Tae-Ho Lee, Hui Guo, Xiyin Wang, Changsoo Kim, and Andrew H Paterson. Snphylo: a pipeline to construct a phylogenetic tree from huge snp data. BMC genomics, 15(1):162, 2014.

34. Wen-Hsiung Li, Masako Tanimura, and Paul M Sharp. An evaluation of the molecular clock hypothesis using mammalian dna sequences. Journal of molecular evolution, 25(4):330–342, 1987.

35. M. Martiniaková, R. Omelka, Jancová, R. Stawarz, and G. Formicki. Heavy metal content in the femora of yellow-necked mouse (apodemus flavicollis) and wood mouse (apodemus sylvaticus) from different types of polluted environment in slovakia. Environmental monitoring and assessment, 171(1-4):651–660, 2010.

36. A Mastretta-Yanes, Nils Arrigo, Nadir Alvarez, Tove H Jorgensen, D Piñero, and BC Emerson. Restriction site-associated dna sequencing, genotyping error estimation and de novo assembly optimization for population genetic inference. Molecular Ecology Resources, 15(1):28–41, 2015.

37. Johan René Michaux, Elodie Magnanou, Emmanuel Paradis, Caroline Nieberding, and Roland Libois. Mitochondrial phylogeography of the woodmouse (apodemus sylvaticus) in the western palearctic region. Molecular Ecology, 12(3):685–697, 2003.

38. JR Michaux, P Chevret, M-G Filippucci, and M Macholan. Phylogeny of the genus apodemus with a special emphasis on the subgenus sylvaemus using the nuclear irbp gene and two mitochondrial markers: cytochrome b and 12s rrna. Molecular phylogenetics and evolution, 23(2):123–136, 2002.

39. JR Michaux, Roland Libois, E Paradis, and M-G Filippucci. Phylogeographic history of the yellow-necked fieldmouse (apodemus flavicollis) in europe and in the near and middle east. Molecular phylogenetics and evolution, 32(3):788–798, 2004.

40. JR Michaux, Roland Libois, and MG Filippucci. So close and so different: comparative phylogeography of two small mammal species, the yellow-necked fieldmouse (apodemus flavicollis) and the woodmouse (apodemus sylvaticus) in the western palearctic region. Heredity, 94(1):52–63, 2005.

41. M.R. Miller, J.P. Dunham, A. Amores, W.A. Cresko, and E.A. Johnson. Rapid and cost-effective polymorphism identification and genotyping using restriction site associated dna (rad) markers. Genome research, 17(2): 240–248, 2007.

42. Luwanika Mlera and Marshall E Bloom. The role of mammalian reservoir hosts in tick-borne flavivirus biology. Frontiers in cellular and infection microbiology, 8:298, 2018.

43. Andre E Moura, John G Kenny, Roy Chaudhuri, Margaret A Hughes, Andreanna J. Welch, Ryan R Reisinger, PJ Nico de Bruyn, Marilyn E Dahlheim, Neil Hall, and A Rus Hoelzel. Population genomics of the killer whale indicates ecotype evolution in sympatry involving both selection and drift. Molecular Ecology, 23(21): 5179–5192, 2014.

44. Eric Pante, Jawad Abdelkrim, Amélia Viricel, Delphine Gey, SC France, Marie-Catherine Boisselier, and Sarah Samadi. Use of rad sequencing for delimiting species. Heredity, 114(5):450, 2015.

45. Anna Papa, Elton Rogozi, Enkelejda Velo, Evangelia Papadimitriou, and Silvia Bino. Genetic detection of hantaviruses in rodents, albania. Journal of medical virology, 88(8):1309–1313, 2016.

46. Josephine R Paris, Jamie R Stevens, and Julian M Catchen. Lost in parameter space: a road map for stacks. Methods in Ecology and Evolution, 2017.

47. Venkatramana Pegadaraju, Rick Nipper, Brent Hulke, Lili Qi, and Quentin Schultz. De novo sequencing of sunflower genome for snp discovery using rad (restriction site associated dna) approach. BMC genomics, 14(1): 556, 2013.

48. Brant K Peterson, Jesse N Weber, Emily H Kay, Heidi S Fisher, and Hopi E Hoekstra. Double digest radseq: an inexpensive method for de novo snp discovery and genotyping in model and non-model species. PloS one, 7(5): e37135, 2012.

49. Jesse A Poland and Trevor W Rife. Genotyping-by-sequencing for plant breeding and genetics. The Plant Genome, 5(3):92–102, 2012.

50. Jesse A Poland, Patrick J Brown, Mark E Sorrells, and Jean-Luc Jannink. Development of high-density genetic maps for barley and wheat using a novel two-enzyme genotyping-by-sequencing approach. PloS one, 7(2): e32253, 2012.

51. Zdzisław Pucek, Jadwiga Stachurska, Magdalena Bibrich, et al. Keys to vertebrates of poland: mammals. 1981.

52. Zdzisław Pucek, Włodzimierz Jędrzejewski, Bogumiła Jędrzejewska, and Michalina Pucek. Rodent population dynamics in a primeval deciduous forest (białowieża national park) in relation to weather, seed crop, and predation. Acta Theriologica, 38(2):199–232, 1993.

53. Sebastien J Puechmaille. The program structure does not reliably recover the correct population structure when sampling is uneven: subsampling and new estimators alleviate the problem. Molecular Ecology Resources, 16(3):608–627, 2016.

54. S Purcell and CC Chang. Plink 1.9 package. 2018. URL https://www.cog-genomics.org/plink/1.9/.

55. Marija Rajičić, Svetlana A Romanenko, Tatyana V Karamysheva, Jelena Blagojević, Tanja Adnađević, Ivana Budinski, Aleksey S Bogdanov, Vladimir A Trifonov, Nikolay B Rubtsov, and Mladen Vujošević. The origin of b chromosomes in yellow-necked mice (apodemus flavicollis)break rules but keep playing the game. PloS one, 12 (3):e0172704, 2017.

56. AM Reitzel, S Herrera, MJ Layden, MQ Martindale, and TM Shank. Going where traditional markers have not gone before: utility of and promise for rad sequencing in marine invertebrate phylogeography and population genomics. Molecular ecology, 22(11):2953–2970, 2013.

57. Dania Richter, Daniela B Schlee, and Franz-Rainer Matuschka. Reservoir competence of various rodents for the lyme disease spirochete borrelia spielmanii. Applied and environmental microbiology, 2011.

58. Adriana Rico, Pavel Kindlmann, and František Sedláček. Can the barrier effect of highways cause genetic subdivision in small mammals? Acta theriologica, 54(4):297–310, 2009.

59. Naiara Rodriguez-Ezpeleta, Paula Álvarez, and Xabier Irigoien. Genetic diversity and connectivity in maurolicus muelleri in the bay of biscay inferred from thousands of snp markers. Frontiers in genetics, 8:195, 2017.

60. Aaron Shafer, Claire R Peart, Sergio Tusso, Inbar Maayan, Alan Brelsford, Christopher W Wheat, and Jochen BW Wolf. Bioinformatic processing of rad-seq data dramatically impacts downstream population genetic inference. Methods in Ecology and Evolution, 8(8):907–917, 2017.

61. Allison J Shultz, Allan J Baker, Geoffrey E Hill, Paul M Nolan, and Scott V Edwards. Snp s across time and space: population genomic signatures of founder events and epizootics in the house finch (haemorhous mexicanus). Ecology and evolution, 6(20):7475–7489, 2016.

62. M. Velickovic. Measures of the developmental stability, body size and body condition in the black-striped mouse (apodemus agrarius) as indicators of a disturbed environment in northern serbia. Belgian Journal of Zoology, 137(2):147, 2007.

63. Mladen Vujoevi, Marija Rajii, and Jelena Blagojevi. B chromosomes in populations of mammals revisited. Genes, 9(10):487, 2018.

