## Supplementary material for "Population structure of Apodemus flavicollis and comparison to Apodemus sylvaticus in northern Poland based on RAD-seq": biorxiv_supp_files.pdf

### Supplementary Material: Population structure of *Apodemus flavicollis* and comparison to *Apodemus sylvaticus* in northern Poland based on whole-genome genotyping with RAD-seq

#### 1 Sample information

Table 1: Sample ID, coordinates and environmental information for the samples. Bory - Bory Tucholskie, Bial - Białowieża, Hack - Haćki, Kadz - Kadzidło

| ID | Species | Location | Latitude | Longitude | Environment |
|---|---|---|---|---|---|
| D04 | A. flavicollis | Bory | 17.58 | 53.77 | mesic pine forest |
| E04 | A. flavicollis | Bory | 17.56 | 53.81 | mesic pine forest |
| F04 | A. flavicollis | Bory | 17.55 | 53.79 | mesic pine forest |
| G04 | A. flavicollis | Bory | 17.58 | 53.78 | dry pine forest |
| H04 | A. flavicollis | Bory | 17.58 | 53.77 | mesic pine forest |
| A05 | A. flavicollis | Bory | 17.58 | 53.77 | mesic pine forest |
| B05 | A. flavicollis | Bory | 17.58 | 53.77 | sedge meadow |
| C05 | A. flavicollis | Bory | 17.58 | 53.77 | mesic pine forest |
| D05 | A. flavicollis | Bory | 17.58 | 53.77 | mesic pine forest |
| E05 | A. flavicollis | Bory | 17.51 | 53.80 | oak-lime-hornbeam forest |
| F05 | A. flavicollis | Bory | 17.58 | 53.77 | mesic pine forest |
| G05 | A. flavicollis | Bory | 17.51 | 53.80 | oak-lime-hornbeam forest |
| H05 | A. flavicollis | Bory | 17.51 | 53.80 | oak-lime-hornbeam forest |
| G12 | A. flavicollis | Bory | 17.51 | 53.80 | oak-lime-hornbeam forest |
| H12 | A. flavicollis | Bory | 17.56 | 53.81 | alder foorest at lake |
| C06 | A. flavicollis | Bory | 17.58 | 53.78 | reeds at lake |

| Sample | Species | Location | Long | Lat | Environment |
|---|---|---|---|---|---|
| D06 | A. flavicollis | Bory | 17.51 | 53.80 | oak-lime-hornbeam forest |
| E06 | A. flavicollis | Bory | 17.51 | 53.80 | oak-lime-hornbeam forest |
| F06 | A. flavicollis | Bory | 17.51 | 53.80 | oak-lime-hornbeam forest |
| H06 | A. flavicollis | Bory | 17.56 | 53.81 | mesic pine forest |
| A07 | A. flavicollis | Bory | 17.56 | 53.81 | alder foorest at lake |
| B07 | A. flavicollis | Bory | 17.58 | 53.78 | mesic pine forest |
| C07 | A. flavicollis | Bory | 17.51 | 53.80 | oak-lime-hornbeam forest |
| A04 | A. flavicollis | Bial | 23.85 | 52.71 | cultivated meadow |
| B04 | A. flavicollis | Bial | 23.85 | 52.71 | cultivated meadow |
| C04 | A. flavicollis | Bial | 23.85 | 52.71 | cultivated meadow |
| F08 | A. flavicollis | Bial | 23.83 | 52.72 | oak-lime-hornbeam forest |
| G08 | A. flavicollis | Bial | 23.83 | 52.72 | oak-lime-hornbeam forest |
| H08 | A. flavicollis | Bial | 23.83 | 52.72 | oak-lime-hornbeam forest |
| A09 | A. flavicollis | Bial | 23.83 | 52.72 | oak-lime-hornbeam forest |
| B09 | A. flavicollis | Bial | 23.83 | 52.72 | oak-lime-hornbeam forest |
| C09 | A. flavicollis | Bial | 23.83 | 52.72 | oak-lime-hornbeam forest |
| D09 | A. flavicollis | Bial | 23.83 | 52.72 | oak-lime-hornbeam forest |
| E09 | A. flavicollis | Bial | 23.83 | 52.72 | oak-lime-hornbeam forest |
| F09 | A. flavicollis | Bial | 23.83 | 52.72 | oak-lime-hornbeam forest |
| G09 | A. flavicollis | Bial | 23.83 | 52.72 | oak-lime-hornbeam forest |
| H09 | A. flavicollis | Bial | 23.83 | 52.72 | oak-lime-hornbeam forest |
| A10 | A. flavicollis | Bial | 23.83 | 52.72 | oak-lime-hornbeam forest |
| B10 | A. flavicollis | Bial | 23.83 | 52.72 | oak-lime-hornbeam forest |
| C10 | A. flavicollis | Bial | 23.83 | 52.72 | oak-lime-hornbeam forest |
| D10 | A. flavicollis | Bial | 23.83 | 52.72 | oak-lime-hornbeam forest |
| E10 | A. flavicollis | Bial | 23.83 | 52.72 | oak-lime-hornbeam forest |
| F10 | A. flavicollis | Bial | 23.85 | 52.72 | oak-lime-hornbeam forest |
| G10 | A. flavicollis | Bial | 23.85 | 52.72 | oak-lime-hornbeam forest |
| H10 | A. flavicollis | Bial | 23.82 | 52.74 | oak-lime-hornbeam forest |
| A11 | A. flavicollis | Bial | 23.82 | 52.75 | oak-lime-hornbeam forest |
| B11 | A. flavicollis | Bial | 23.82 | 52.70 | sedge meadow |
| C11 | A. flavicollis | Bial | 23.85 | 52.72 | oak-lime-hornbeam forest |
| D11 | A. flavicollis | Bial | 23.85 | 52.72 | oak-lime-hornbeam forest |
| E11 | A. flavicollis | Bial | 23.85 | 52.72 | oak-lime-hornbeam forest |
| F11 | A. flavicollis | Bial | 23.85 | 52.72 | oak-lime-hornbeam forest |

| Sample | Species | Location | Long | Lat | Environment |
|---|---|---|---|---|---|
| G11 | A. flavicollis | Bial | 23.85 | 52.72 | oak-lime-hornbeam forest |
| H11 | A. flavicollis | Bial | 23.85 | 52.72 | oak-lime-hornbeam forest |
| A12 | A. flavicollis | Bial | 23.85 | 52.72 | oak-lime-hornbeam forest |
| B12 | A. flavicollis | Bial | 23.85 | 52.72 | oak-lime-hornbeam forest |
| C12 | A. flavicollis | Bial | 23.85 | 52.72 | oak-lime-hornbeam forest |
| D12 | A. flavicollis | Bial | 23.85 | 52.72 | oak-lime-hornbeam forest |
| E12 | A. flavicollis | Bial | 23.85 | 52.72 | oak-lime-hornbeam forest |
| C02 | A. flavicollis | Hack | 23.17 | 52.83 | xerothermic meadow |
| D02 | A. flavicollis | Hack | 23.17 | 52.83 | xerothermic meadow |
| E02 | A. flavicollis | Hack | 23.17 | 52.83 | xerothermic meadow |
| F02 | A. flavicollis | Hack | 23.17 | 52.83 | xerothermic meadow |
| G02 | A. flavicollis | Hack | 23.17 | 52.83 | xerothermic meadow |
| H02 | A. flavicollis | Hack | 23.17 | 52.83 | xerothermic meadow |
| A03 | A. flavicollis | Hack | 23.17 | 52.83 | xerothermic meadow |
| B03 | A. flavicollis | Hack | 23.17 | 52.83 | xerothermic meadow |
| C03 | A. flavicollis | Hack | 23.17 | 52.83 | xerothermic meadow |
| D03 | A. flavicollis | Hack | 23.17 | 52.83 | xerothermic meadow |
| E03 | A. flavicollis | Hack | 23.17 | 52.83 | xerothermic meadow |
| F03 | A. flavicollis | Hack | 23.17 | 52.83 | xerothermic meadow |
| G03 | A. flavicollis | Hack | 23.17 | 52.83 | xerothermic meadow |
| H03 | A. flavicollis | Hack | 23.17 | 52.83 | xerothermic meadow |
| D07 | A. sylvaticus | Bory | 17.54 | 53.79 | dry pine forest |
| E07 | A. sylvaticus | Bory | 17.56 | 53.79 | reeds at lake |
| F07 | A. sylvaticus | Bory | 17.54 | 53.79 | dry pine forest |
| G07 | A. sylvaticus | Bory | 17.55 | 53.79 | mesic pine forest |
| H07 | A. sylvaticus | Bory | 17.54 | 53.79 | dry pine forest |
| A08 | A. sylvaticus | Kadz | 21.37 | 53.20 | dry pine forest |
| B08 | A. sylvaticus | Kadz | 21.37 | 53.20 | dry pine forest |
| C08 | A. sylvaticus | Kadz | 21.37 | 53.20 | dry pine forest |
| D08 | A. sylvaticus | Kadz | 21.37 | 53.20 | dry pine forest |
| E08 | A. sylvaticus | Kadz | 21.37 | 53.20 | dry pine forest |

### 2 Barcodes and demultiplexing

Table 2: Barcodes used and demultiplexing results.

| Barcode | Filename | Total | NoRadTag | LowQuality | Retained |
|---|---|---|---|---|---|
| TATTCGCAT | D01 | 1754624 | 823059 | 322 | 802448 |
| CCTTGCCATT | B02 | 5616138 | 2656224 | 1136 | 2524650 |
| GGTATA | C02 | 2497516 | 1178292 | 412 | 1112807 |
| TCTTGG | D02 | 1925168 | 926463 | 359 | 862961 |
| GGTGT | E02 | 1772988 | 836727 | 301 | 802310 |
| GGATA | F02 | 2103232 | 990036 | 342 | 950445 |
| CTAAGCA | G02 | 2396754 | 1118652 | 420 | 1107411 |
| ATTAT | H02 | 3592492 | 1672167 | 603 | 1654087 |
| GCGCTCA | A03 | 1701066 | 796055 | 316 | 766110 |
| ACTGCGAT | B03 | 2859122 | 1379563 | 513 | 1244319 |
| TTCGTT | C03 | 2522570 | 1203310 | 467 | 1127262 |
| ATATAA | D03 | 1448256 | 675350 | 261 | 664835 |
| TGGCAACAGA | E03 | 1907170 | 896741 | 415 | 854373 |
| CTCGTCG | F03 | 1424136 | 661282 | 253 | 647835 |
| GCCTACCT | G03 | 1316424 | 631751 | 267 | 579215 |
| CACCA | H03 | 4119158 | 1904665 | 717 | 1918252 |
| AATTAG | A04 | 3353928 | 1576668 | 531 | 1528167 |
| GGAACGA | B04 | 2714032 | 1268460 | 499 | 1237675 |
| ACTGCT | C04 | 1519814 | 732180 | 279 | 676595 |
| TGCTT | D04 | 3337318 | 1598963 | 538 | 1516482 |
| GCAAGCCAT | E04 | 2272530 | 1077974 | 436 | 1028556 |
| CGCACCAATT | F04 | 1328064 | 629634 | 257 | 597209 |
| CTCGCGG | G04 | 2843128 | 1352618 | 497 | 1300936 |
| AACTGG | H04 | 1773388 | 851270 | 311 | 799274 |
| ATGAGCAA | A05 | 3543298 | 1701957 | 692 | 1580365 |
| CTTGA | B05 | 2280988 | 1099255 | 413 | 1016552 |
| GCGTCCT | C05 | 3835930 | 1834408 | 674 | 1724309 |
| ACCAGGA | D05 | 3081248 | 1488008 | 581 | 1378175 |
| CCACTCA | E05 | 2003682 | 940201 | 332 | 919846 |
| TCACGGAAG | F05 | 889424 | 420138 | 187 | 407176 |
| TATCA | G05 | 1212906 | 593550 | 171 | 545872 |
| TAGCCAA | H05 | 1794800 | 838457 | 312 | 836413 |

| Barcode | Filename | Total | NoRadTag | LowQuality | Retained |
|---|---|---|---|---|---|
| GGTGCACATT | C06 | 1784198 | 845365 | 349 | 798706 |
| CTCTCGCAT | D06 | 1495486 | 710272 | 290 | 675749 |
| CAGAGGT | E06 | 1827948 | 891153 | 317 | 810051 |
| GCGTACAAT | F06 | 1083614 | 509870 | 219 | 494520 |
| ACGCGCG | G06 | 1490100 | 697737 | 247 | 686381 |
| GTCGCCT | H06 | 2562952 | 1219598 | 434 | 1168312 |
| AATAACCAA | A07 | 2750168 | 1290585 | 509 | 1254852 |
| AATGAACGA | B07 | 2023934 | 970073 | 414 | 904065 |
| ATGGCAA | C07 | 2897680 | 1386103 | 501 | 1307711 |
| GAAGCA | D07 | 4523918 | 2130884 | 804 | 2088074 |
| AACGTGCCT | E07 | 3561580 | 1678483 | 705 | 1636000 |
| CCTCG | F07 | 4775646 | 2243939 | 795 | 2212819 |
| CTCAT | G07 | 2816492 | 1342690 | 453 | 1290660 |
| ACGGTACT | H07 | 1538254 | 721448 | 267 | 710639 |
| GCGCCG | A08 | 1581588 | 751487 | 306 | 715924 |
| CAAGT | B08 | 2362354 | 1126323 | 386 | 1076036 |
| GGAGTCAAG | C08 | 1931910 | 921128 | 340 | 866712 |
| TGAAT | D08 | 2004632 | 977907 | 329 | 903723 |
| CATAT | E08 | 2845620 | 1348639 | 463 | 1306750 |
| GTGACACAT | F08 | 1793344 | 840977 | 320 | 798641 |
| TATGT | G08 | 1912488 | 889366 | 326 | 892469 |
| TGCAGA | H08 | 1587072 | 744356 | 247 | 728265 |
| CATCTGCCG | A09 | 1927106 | 894811 | 400 | 865134 |
| GGACAG | B09 | 2391890 | 1139966 | 395 | 1084539 |
| ATCTGT | C09 | 4006790 | 1882256 | 717 | 1829175 |
| AAGACGCT | D09 | 2083594 | 1008686 | 376 | 912137 |
| GAATGCAATA | E09 | 1673516 | 809802 | 316 | 720740 |
| TAGCAG | F09 | 1611016 | 772260 | 268 | 720206 |
| CTTAG | G09 | 1236082 | 639672 | 195 | 503807 |
| TTATTACAT | H09 | 903066 | 480838 | 156 | 346810 |
| GCCAACAAGA | A10 | 2280156 | 1095405 | 452 | 1002024 |
| TGCCGCAT | B10 | 4328430 | 2092480 | 779 | 1906904 |
| CGTGTCA | C10 | 2174200 | 1069375 | 366 | 944630 |
| CAACCACACA | D10 | 1994002 | 989286 | 367 | 844802 |
| GCTCCGA | E10 | 2269544 | 1072489 | 435 | 1027809 |

| Barcode | Filename | Total | NoRadTag | LowQuality | Retained |
|---|---|---|---|---|---|
| CGTTCA | F10 | 2396728 | 1134028 | 402 | 1063780 |
| CATCACAAG | G10 | 1130460 | 547376 | 213 | 482972 |
| TCCAG | H10 | 1134466 | 543467 | 172 | 500967 |
| AACTGAAG | A11 | 2060310 | 972421 | 383 | 912133 |
| GATTCA | B11 | 1559246 | 726131 | 226 | 712556 |
| CAAGCCAATT | C11 | 2759210 | 1374653 | 491 | 1161521 |
| TTGCGCT | D11 | 2013912 | 958443 | 341 | 907508 |
| CGCAGACACT | E11 | 1773052 | 884978 | 334 | 742494 |
| TGTGGA | F11 | 1638142 | 778419 | 287 | 738748 |
| TGGATA | G11 | 2001520 | 982084 | 343 | 878640 |
| ATAGCGT | H11 | 1929208 | 896385 | 353 | 888463 |
| CCATAGA | A12 | 4910032 | 2303281 | 844 | 2177377 |
| GGCACGCAT | B12 | 5959610 | 2968959 | 1184 | 2473214 |
| ATTAACAATT | C12 | 1040872 | 496149 | 188 | 452788 |
| CAATA | D12 | 2431454 | 1185404 | 393 | 1053547 |
| TAGTCCAT | E12 | 1326952 | 615885 | 252 | 615685 |
| CGTGACCT | F12 | 1389748 | 657705 | 274 | 614681 |
| CTTCAGA | G12 | 9249428 | 4445105 | 1627 | 4157586 |
| ATCTGCAACA | H12 | 3268002 | 1557924 | 623 | 1457762 |
| | **total** | **206744014** | **98568584** | **36987** | **92741120** |
| | **min** | **889424** | **420138** | **156** | **346810** |
| | **max** | **9249428** | **4445105** | **1627** | **4157586** |
| | **average** | **2404000.16** | **1146146.32** | **430.08** | **1078385.11** |
| | **stdev** | **1282551.13** | **613923.28** | **234.71** | **575871.53** |
| | **median** | **2009272** | **975164** | **366.5** | **905786.5** |

### 3 Estimation of the best parameters for the combined dataset

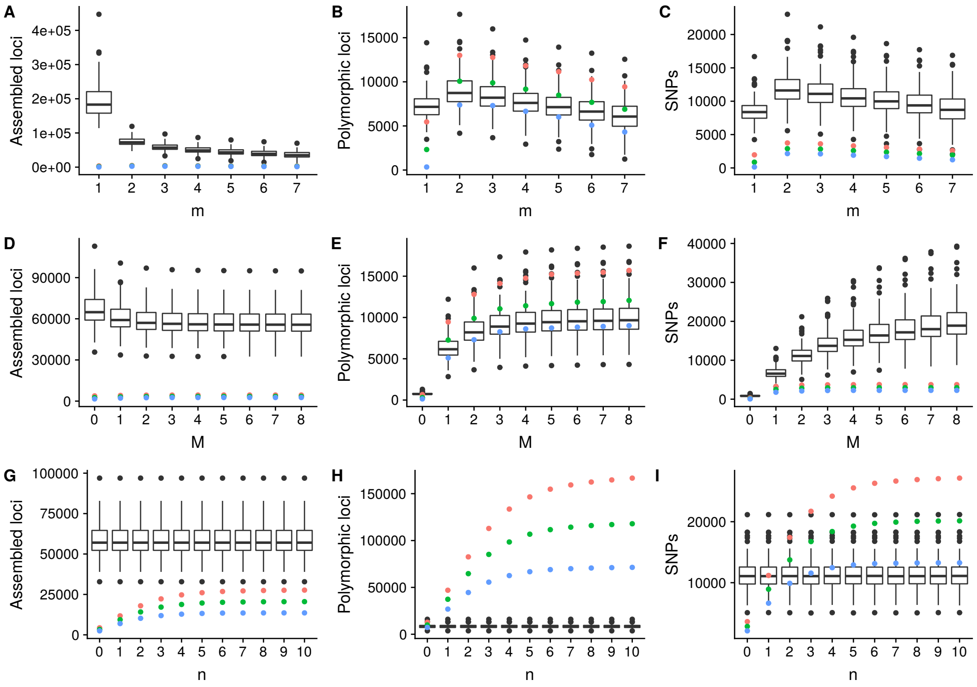


Figure 1: Selection of the optimal parameters for the combined dataset. Number of assembled loci, polymorphic loci and SNPs for iterating values of m, M and n parameters. Blue circles represent data found in at least 40% of the population, green circles in the 60% and red circles in the 80%. Data used to build the figure is available on GitHub: `https://github.com/Marisa89/ddRADseq_poland/blob/master/Tables/Apodemus/Table_selection_best_parameters_Apodemus.xlsx`

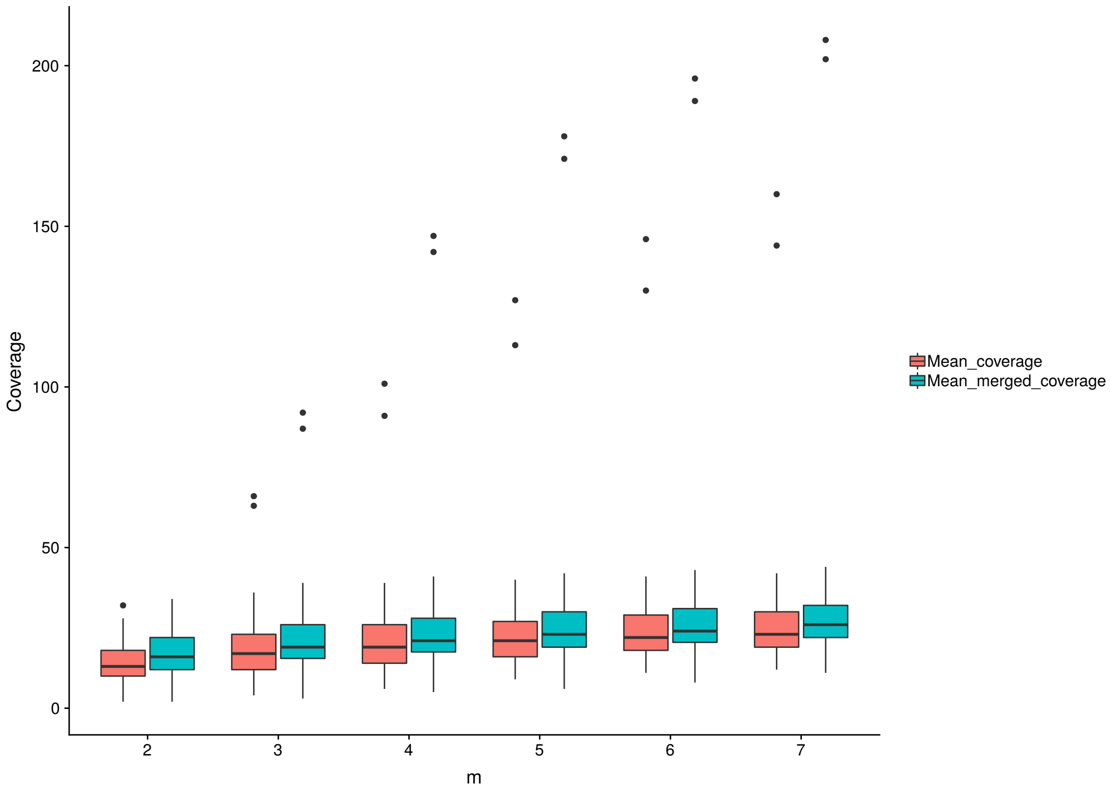


Figure 2: .

Distribution of the mean coverage before and after merging loci for each iteration of the m parameter.Results for the combined dataset including samples from *Apodemus flavicollis* and *Apodemus sylvaticus*. Data used to build the figure is available on GitHub: `https://github.com/Marisa89/ddRADseq_poland/blob/master/Tables/Apodemus/Table_coverage_Apodemus.csv`

### 4 Estimation of the best parameters for *Apodemus flavicollis* samples

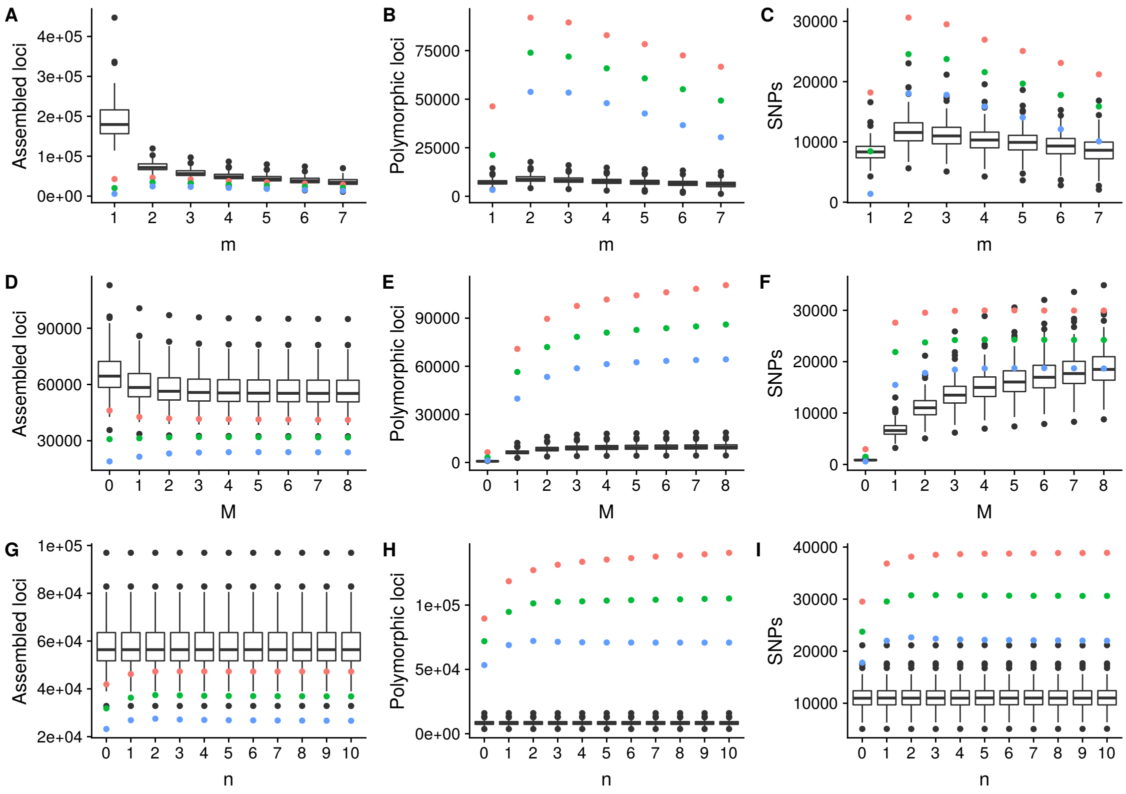


Figure 3: Number of assembled loci, polymorphic loci and SNPs for iterating values of m, M and n parameters. Blue circles represent data found in at least 40% of the individuals, green circles in the 60% and red circles in the 80%. Data used to build the figure is available on GitHub: `https://github.com/Marisa89/ddRADseq_poland/blob/master/Tables/A.flavicollis/Table_selection_best_parameters_Aflavicollis.xlsx`

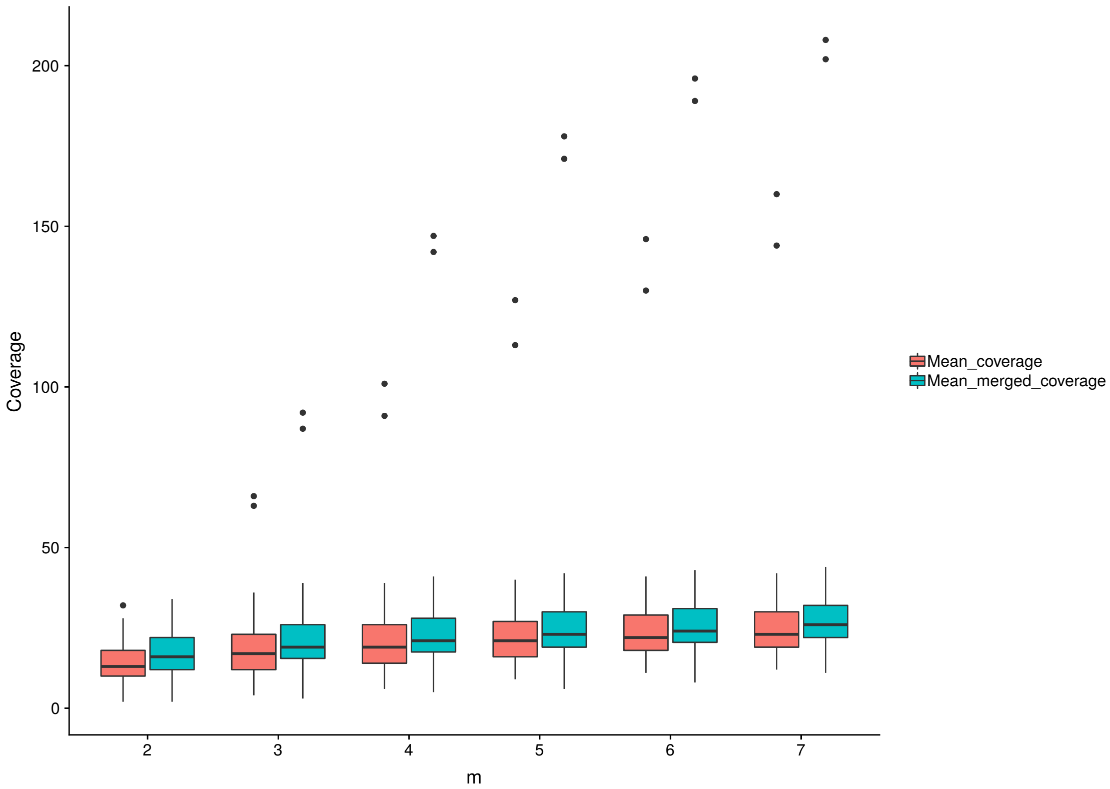


Figure 4: Distribution of the mean coverage before and after merging loci for each iteration of the m parameter for *Apodemus flavicollis* samples. Data used to build the figure is available on GitHub: `https://github.com/Marisa89/ddRADseq_poland/blob/master/Tables/A.flavicollis/Table_coverage_Aflavicollis.csv`

#### 5 Cross-validation errors

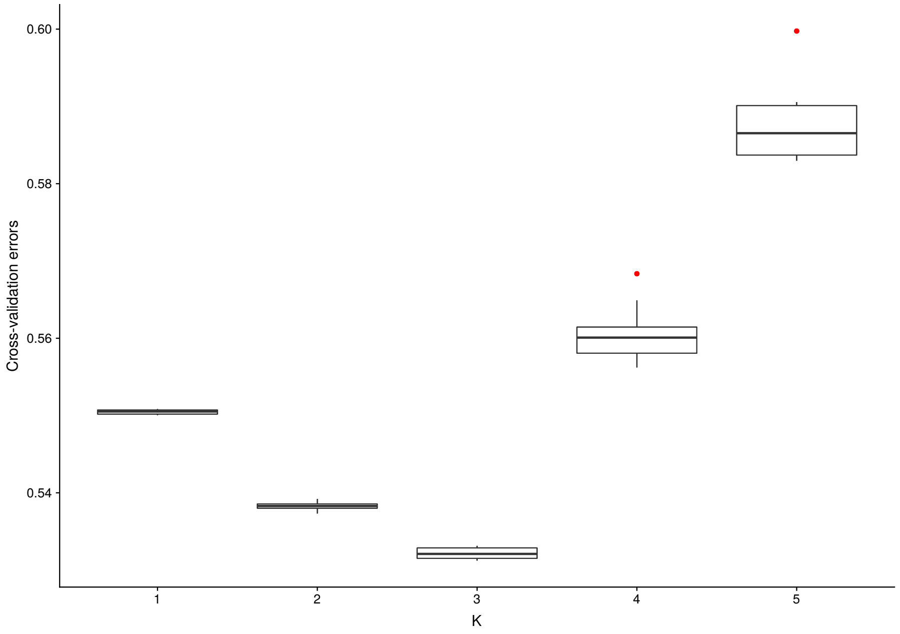


Figure 5: Cross-validation errors obtained for values of K between 1 and 5 for 10 runs with different seeds for all samples

#### 6 Catalogue of loci used for species differentiation

Due to the size of the catalogue, the files has been uploaded into Dropbox. They are available for download at the following link: `https://www.dropbox.com/sh/3757wzer94eef85/AADRXN6GT5J6QJ-JHFySi34Aa?dl=0`

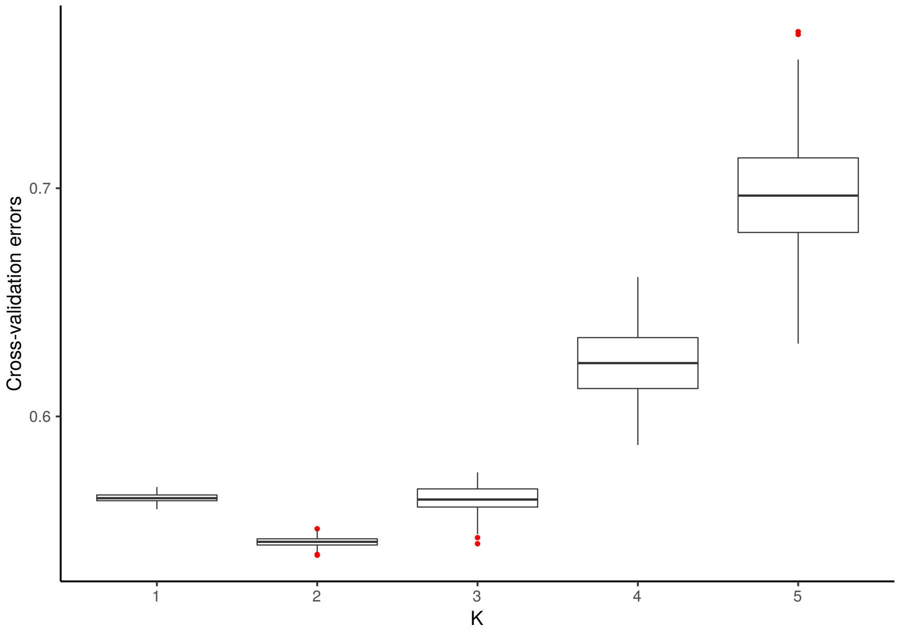


Figure 6: Cross-validation errors obtained for values of K between 1 and 5 for the 100 permutations performed with randomly-drawn equal number of samples per population (n = 15)

### 7 117 loci with the highest divergence

Table 3: List of 117 loci with the highest divergence between both species

| **List of loci with a divergence higher than 4.9%** | | | | | |
|---|---|---|---|---|---|
| 3211 | 11103 | 20032 | 35338 | 45028 | 62435 |
| 4189 | 11112 | 20410 | 35417 | 45908 | 62495 |
| 4759 | 12321 | 20426 | 35799 | 47463 | 62544 |
| 4835 | 12823 | 21475 | 36256 | 51367 | 62719 |
| 4967 | 13690 | 22268 | 36342 | 51435 | 62846 |
| 5241 | 13708 | 23146 | 36597 | 51533 | 64055 |
| 5937 | 13820 | 23682 | 36821 | 53072 | 64057 |
| 6024 | 14596 | 24277 | 37171 | 53520 | 64228 |
| 6497 | 14916 | 25086 | 37193 | 53551 | 64457 |
| 6678 | 15177 | 25874 | 38518 | 53831 | 64631 |
| 7484 | 15553 | 26440 | 39788 | 54014 | 65038 |
| 7873 | 16614 | 26520 | 39844 | 57051 | 65147 |
| 8108 | 16806 | 27415 | 39936 | 57466 | 65161 |
| 8225 | 17192 | 30030 | 40266 | 59850 | 65163 |
| 9035 | 17594 | 31857 | 40440 | 60100 | 66267 |
| 9762 | 18137 | 32033 | 41161 | 60367 | 66602 |
| 10097 | 18207 | 32483 | 42388 | 60452 | 67679 |
| 10594 | 19036 | 32926 | 42581 | 61260 | |
| 10967 | 19729 | 33371 | 42639 | 61310 | |
| 11041 | 19799 | 33510 | 42900 | 62087 | |

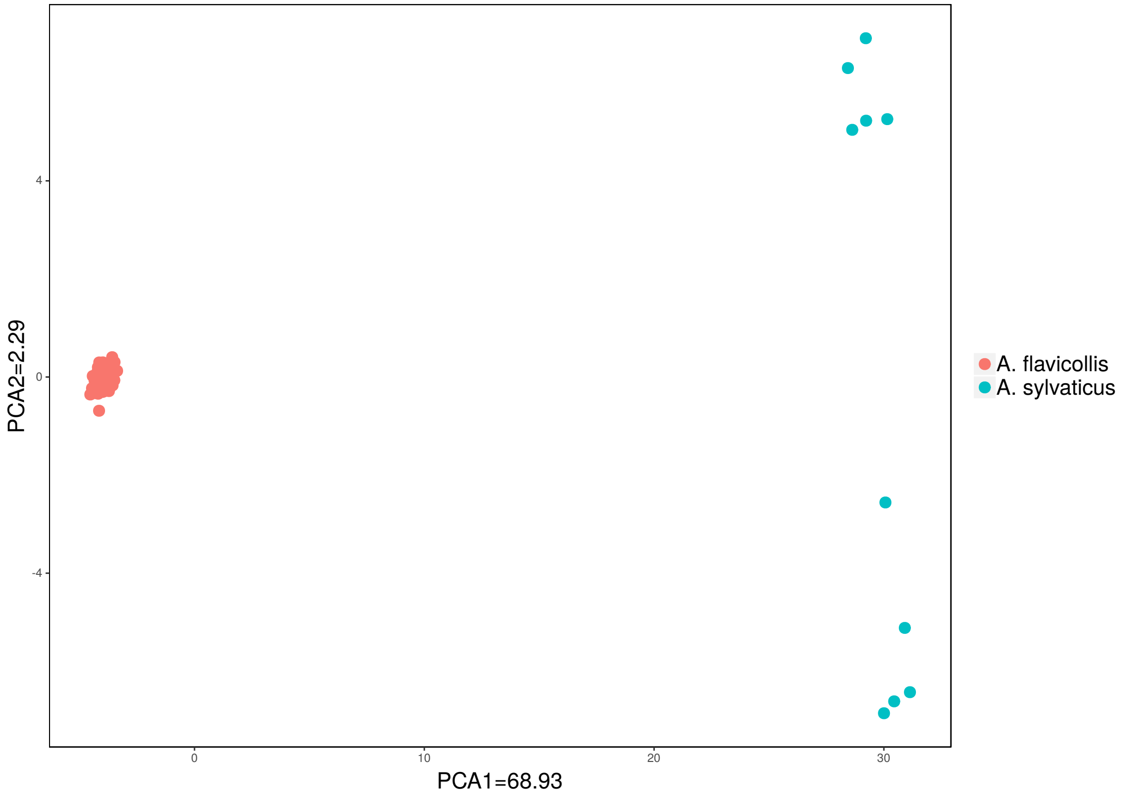


Figure 7: Principal Component Analysis using only the 117 loci with the highest divergence to differentiate Polish samples of *A. flavicollis* and *A. sylvaticus*

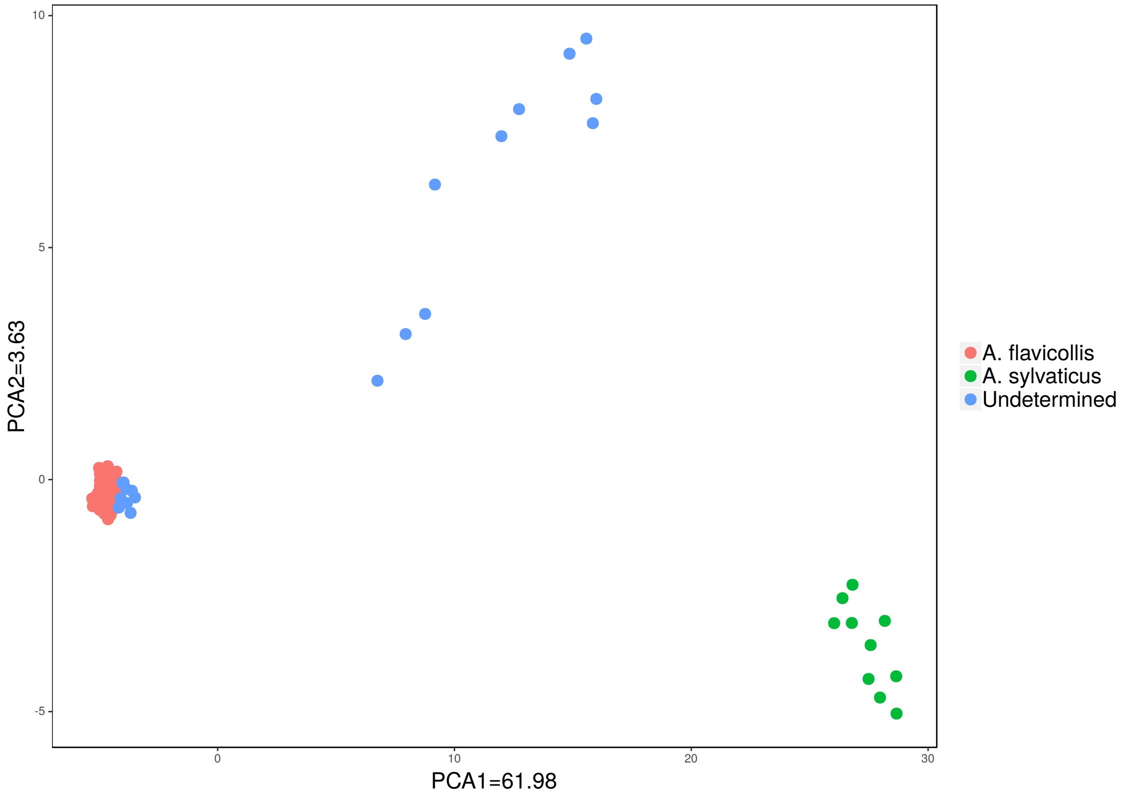


Figure 8: Principal Component Analysis using only the 117 loci with the highest divergence for the dataset including all Polish as well as other European and Tunisian samples.

#### 8 European and Tunisian samples

Table 4: Details of the 20 European and Tunisian samples used to check the catalogue of loci generated based on the Polish samples.

| **ID** | **Source** | **Code** |
|---|---|---|
| AT1 | Johan Michaux | JRM-203 |
| AT2 | Johan Michaux | JRM-204 |
| LT1 | Karol Zub | JB-466 |
| LT2 | Karol Zub | JB-468 |
| LT3 | Karol Zub | JB-470 |
| LT4 | Karol Zub | JB-485 |
| LT5 | Karol Zub | JB-475 |
| RO1 | Johan Michaux | JRM-2729 |
| RO2 | Johan Michaux | JRM-2720 |
| RO3 | Johan Michaux | JRM-2721 |
| WL1 | National Museums Scotland | NMS.Z.2009.101.1295M |
| WL2 | National Museums Scotland | NMS.Z.2009.101.1296M |
| WL3 | National Museums Scotland | NMS.Z.2009.101.1203M |
| WL4 | National Museums Scotland | NMS.Z.2009.101.1294M |
| TN1 | Johan Michaux | JRM-138 |
| TN2 | Johan Michaux | JRM-139 |
| TN3 | Johan Michaux | JRM-140 |
| SC1 | National Museums Scotland | NMS.Z.2009.101.1M |
| SC2 | National Museums Scotland | NMS.Z.2009.101.2M |
| SC3 | National Museums Scotland | NMS.Z.2009.101.3M |

### 9 Code

Scripts are available in the following repositories at GitHub:

`https://github.com/Marisa89/ddRADseq_poland/tree/master/Code` 1- Demultiplex_concatenation.sh
2- Iteration_parameter_selection.sh
3- Graphs_Iteration_parameters.R
4- PCA_plots_species.R
5- PCA_plots_flavicollis.R
6- Generate_files_for_divergence.sh
7- SNP_error_rate.sh
8- Loci_Allele_error_rate.sh
9- Allele_error_rate.R
10- Permutations.sh
11- Permutations_genetic_diversity_and_Fst_tables.R
12- Admixture_different_seed.sh

The code used to calculate divergence is available at: `https://github.com/jarekbryk/divergenceR`
